# UBAP2L drives scaffold assembly of nuclear pore complexes at the intact nuclear envelope

**DOI:** 10.1101/2023.08.21.554160

**Authors:** Yongrong Liao, Leonid Andronov, Xiaotian Liu, Junyan Lin, Lucile Guerber, Linjie Lu, Arantxa Agote-Arán, Evanthia Pangou, Li Ran, Charlotte Kleiss, Mengdi Qu, Stephane Schmucker, Luca Cirillo, Zhirong Zhang, Daniel Riveline, Monica Gotta, Bruno P. Klaholz, Izabela Sumara

## Abstract

Assembly of macromolecular complexes at correct cellular sites is crucial for cell function. Nuclear pore complexes (NPCs) are large cylindrical assemblies with eightfold rotational symmetry, built through hierarchical binding of nucleoporins (Nups) forming distinct subcomplexes. Here, we uncover a direct role of ubiquitin-associated protein 2-like (UBAP2L) in the biogenesis of properly organized and functional NPCs at the intact nuclear envelope (NE) in human cells. UBAP2L localizes to the nuclear pores and drives the formation of the Y-complex, an essential scaffold component of the NPC, and its localization to the NE. UBAP2L facilitates the interaction of the Y-complex with POM121 and Nup153, the critical upstream factors in a well-defined sequential order of Nups assembly onto NE during interphase. Timely localization of the cytoplasmic Nup transport factor fragile X-related protein 1 (FXR1) to the NE and its interaction with the Y-complex are likewise dependent on UBAP2L. Thus, this NPC biogenesis mechanism integrates the cytoplasmic and the nuclear NPC assembly signals and ensures efficient nuclear transport, adaptation to nutrient stress and cellular proliferative capacity, highlighting the importance of NPC homeostasis at the intact nuclear envelope.

**Teaser:** Liao et al. show how UBAP2L drives the assembly of the scaffold elements into symmetrical and functional NPCs at the nuclear envelope in human cells.

## Introduction

Nuclear pore complexes (NPCs) are among the largest and the most intricate multiprotein assemblies in eukaryotic cells. They constitute the sole communication gates between the nucleus and the cytoplasm thereby ensuring cellular function and survival. NPCs are inserted in the nuclear envelope (NE), a double membrane structure surrounding the cell nucleus, and mediate the transport of proteins and RNAs between the two cellular compartments (Hampoelz *et al*, 2019; Knockenhauer & Schwartz, 2016). Multiple copies of around 30 different nucleoporins (Nups) are the building protein units of the NPCs. Nups initially form various sub-complexes which can subsequently co-assemble, following a hierarchical principle, into functional NPCs (Onischenko *et al*, 2020). The mature NPCs contain a scaffold that surrounds and anchors the Nups with disordered domains forming the inner passage channel (so called Phenylalanine-Glycine repeat Nups or FG-Nups), as well as two asymmetric complex components, the cytoplasmic filaments facing the cytoplasmic side of the NE and the nuclear basket pointing towards the inside of the nucleus. How these architectural elements of the NPC are assembled at the intact NE represents an intriguing and unresolved biological question.

Previous studies using biochemical and high-resolution structural techniques revealed the eightfold rotational symmetry as a characteristic feature of the NPC three-dimensional organization (Beck & Hurt, 2017; Grossman *et al*, 2012; Hampoelz *et al*, 2019; Knockenhauer & Schwartz, 2016; Lin & Hoelz, 2019). One of the main components of the NPC scaffold is the evolutionarily conserved Y-complex (also known as Nup107-160 complex) forming the cytoplasmic and the nuclear rings that encompass the inner ring of the NPC (von Appen *et al*, 2015). In metazoans, the Y-complex is composed of Nup133, Nup107, Nup96 and Sec13, Nup160, Nup37, Elys, Nup85, Seh1 (also named Seh1l) and Nup43 and it is critical for NPC assembly (Doucet *et al*, 2010; Walther *et al*, 2003). Interestingly, FG-Nups can also build the links with the structural scaffold elements and contribute to the biogenesis of the NPC in yeast (Onischenko *et al*, 2017). In metazoan cells, NPCs are formed concomitantly with the reassembly of the NE during mitotic exit but the interphase pathway also exists where NPCs can be formed *de novo* and are inserted into the intact NE through an inside-out mechanism (Otsuka *et al*, 2016). Nup153 and POM121 are the critical upstream components in a well-defined sequential order of Nups assembly onto the interphase nuclei (Otsuka *et al*, 2016; Weberruss & Antonin, 2016). In addition, fragile X-related protein 1 (FXR1) can interact with cytoplasmic Y-complex Nups and facilitate their localization to the NE during interphase through a microtubule-and dynein-dependent mechanism, contributing to the NPC homeostasis (Agote-Aran *et al*, 2020; Agote-Arán *et al*, 2021; Holzer & Antonin, 2020).

However, the crosstalk between the nuclear (POM121, Nup153) and the cytoplasmic (FXR1) determinants of the NPC assembly during interphase and the pathways governing the formation of the essential NPC sub-complexes (such as the Y-complex) at the intact NE, remained unexplored. Likewise, it remained unknown what are the signaling pathways defining the oligomerization state of these scaffold elements and ultimately the assembly of the eightfold-symmetrical NPC. Here, we uncover a molecular mechanism based on UBAP2L protein which links the cytoplasmic and the nuclear NPC assembly signals and by which human cells can build the scaffold elements into functional NPCs at the NE during interphase, thereby ensuring cellular function and survival.

## Results

### UBAP2L localizes to the NPCs and interacts with Nups and NPC assembly factors

NPC assembly during interphase is particularly active as cells grow during early G1 phase where an increase in NPC biogenesis has been observed immediately after NE reformation (Dultz & Ellenberg, 2010; Rampello *et al*, 2020). The number of NPCs can be also modulated in response to cellular needs, for instance during differentiation processes or in carcinogenesis when the density of NPCs and nucleocytoplasmic trafficking augment dramatically (Kau *et al*, 2004). UBAP2L (also known as NICE-4) has been associated with the development of various types of cancer (Chai *et al*, 2016; He *et al*, 2018; Li & Huang, 2014; Ye *et al*, 2017; Zhao *et al*, 2015; Guerber *et al*, 2022), however, the cellular mechanisms underlying its oncogenic potential remain currently unknown. In search for additional biological functions of UBAP2L, we analyzed its subcellular localization using immunofluorescence microscopy and the antibody specifically recognizing endogenous UBAP2L protein. Consistent with published findings (Cirillo *et al*, 2020; Youn *et al*, 2018; Huang *et al*, 2020a; Maeda *et al*, 2016), UBAP2L localized to stress granules (SGs) upon exposure to stress by sodium arsenite, but a weak UBAP2L signal was also found in the nucleus (Fig. 1A) as demonstrated previously (Asano-Inami *et al*, 2023). Likewise, in cells not treated with sodium arsenite, we observed a fraction of endogenous (Fig. 1A) as well as ectopically expressed GFP- (Fig. 1B) and Flag-tag (Fig. 1C) UBAP2L protein to be localized at the NE during interphase. Moreover, UBAP2L was able to accumulate in the nucleus upon treatment with the Leptomycin B (inhibitor of nuclear export factor Exportin 1) similar to the dual specificity protein kinase MPS1 (also known as TTK) which is known to shuttle between nucleus and cytoplasm in interphase cells (Jia *et al*, 2015) (Fig. S1, A to C). These results indicate that UBAP2L also shuttles between these two compartments. Cellular fractionation experiments and western blotting confirmed that a fraction of UBAP2L could be found in the nucleus in interphase (Fig. S1D), in accordance with our published findings (Guerber *et al*, 2023). NE localization of endogenous UBAP2L was detected in early prophase, late telophase and in G1 cells (Fig. S2A), suggesting the role of this protein at the sealed nuclear envelope.

**Fig. 1.**
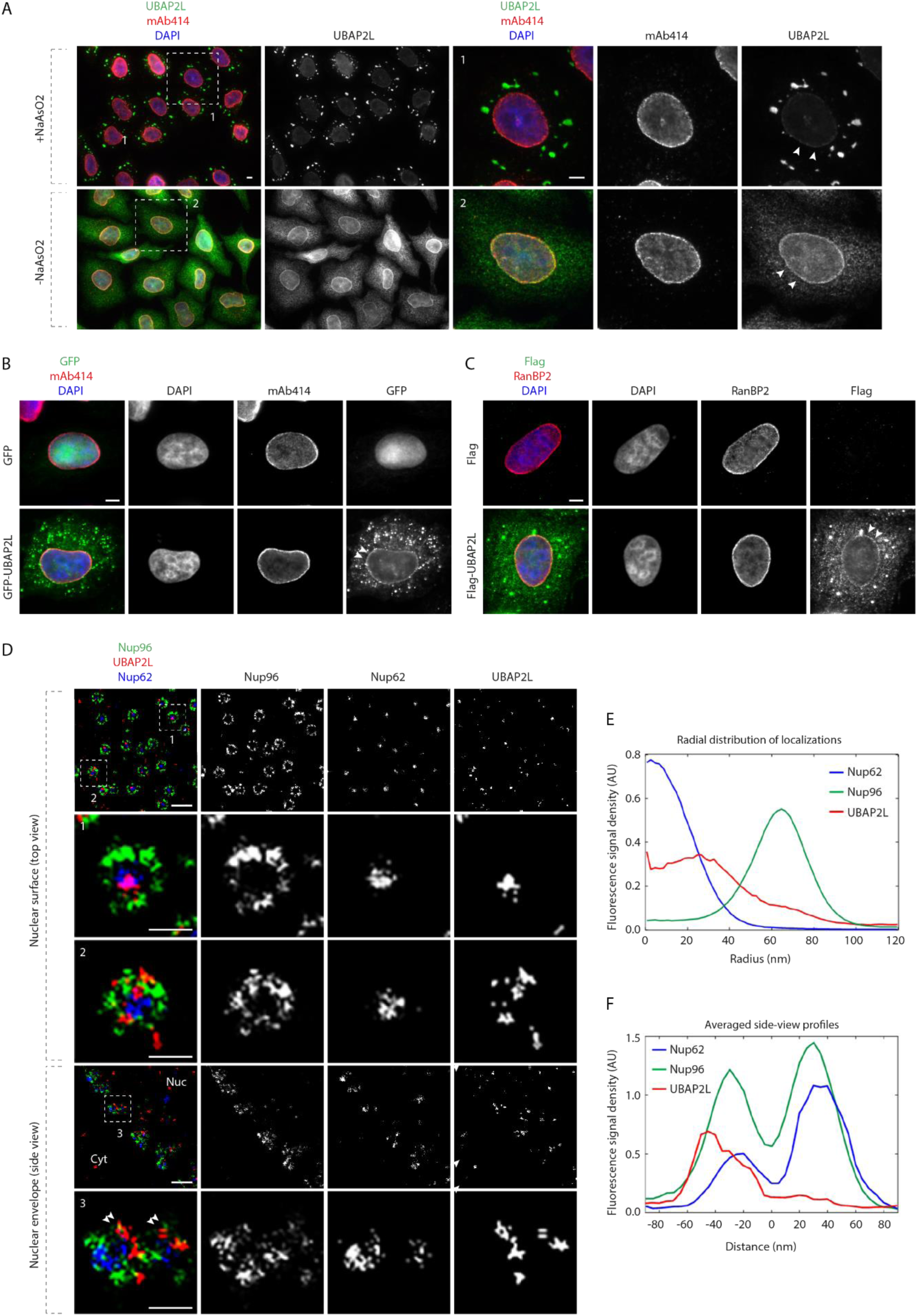
UBAP2L localizes to the NE and NPCs. **(A)** Representative images of the localization of UBAP2L and nucleoporins (Nups) in HeLa cells with/without NaAsO2 treatment shown by immunofluorescence microscopy with UBAP2L and mAb414 antibodies. Nuclei were stained with DAPI. The arrowheads indicate the nuclear envelope (NE) localization of endogenous UBAP2L. The magnified framed regions are shown in the corresponding numbered panels. Scale bars, 5 μm. **(B)** Representative immunofluorescence images depicting the localization of UBAP2L and Nups (mAb414) in HeLa cells expressing GFP alone or GFP-UBAP2L. The arrowheads indicate the NE localization of GFP-tagged UBAP2L. Scale bar, 5 μm. **(C)** Representative immunofluorescence images depicting the localization of UBAP2L and Nups (RanBP2) in HeLa cells expressing Flag alone or Flag-UBAP2L. The arrowheads indicate the NE localization of Flag-tagged UBAP2L. Scale bar, 5 μm (**D to F**) Representative super-resolution immunofluorescence images of Nup96-GFP Knock-in (KI) U2OS cells acquired using multi-color single molecule localization microscopy with a dichroic image splitter (splitSMLM) show NPCs on the nuclear surface (top view) and in the cross section of the NE (side view). Nup96 signal labels the cytoplasmic and nuclear ring of the NPC and the localization of the central channel NPC component is analyzed by Nup62 antibody. Nuclear (Nuc) and cytoplasmic (Cyt) side of the NE are indicated in the side view. The magnified framed regions are shown in the corresponding numbered panels. Note that UBAP2L can localize to both structures within the NPCs (framed regions 1 and 2 in the top view) and is found preferentially at the nuclear ring labelled with Nup96 (double arrowheads in framed region 3 in the side view). Scale bars, 300 and 100 nm, respectively (**D**). Radial distribution of localizations of Nup62, Nup96 and UBAP2L in (**D**) was obtained by averaging 1932 NPC particles (**E**). Averaged “side view” profiles of Nup62, Nup96 and UBAP2L in (**D**) were obtained by alignment of 83 individual NPCs (**F**).

To dissect the nuclear UBAP2L localization more precisely, we used multi-color ratiometric single molecule localization microscopy with a dichroic image splitter (splitSMLM) analysis (Andronov *et al*, 2022, 2021). The splitSMLM analysis revealed that UBAP2L is localized at the NPCs embedded in the NE, where it was found both in the central channel labelled by Nup62 and surrounding the nuclear and cytoplasmic rings labelled by Nup96 of the NPCs (Fig. 1, D to F). Interestingly, fluorescence intensity quantifications indicated that UBAP2L is frequently localized at the side of the Nup96-positive nuclear ring (Fig. 1F). Given that the used super-resolution technique makes it possible to obtain fluorescence images with a resolution in the 20 nm range (Andronov *et al*, 2022), our results suggest that UBAP2L co-localizes with several Nups and building elements of the NPCs at the NE.

These observations prompted us to analyze any possible interactions of UBAP2L with the Nups and the NPC-assembling factors. As expected, immunoprecipitations (IPs) of ectopically expressed GFP-Nup85 in HeLa cells demonstrated an interaction with endogenous Y-complex Nups Nup133 and SEC13 (Doucet *et al*, 2010; Walther *et al*, 2003), with POM121 and Nup153, responsible for targeting Y-complexes to the NE (Otsuka *et al*, 2016; Weberruss & Antonin, 2016) and with the cytoplasmic Nup transporter FXR1 (Agote-Aran *et al*, 2020). GFP-Nup85 also co-immunoprecipitated with endogenous UBAP2L in this analysis (Fig. 2A). In addition, endogenous UBAP2L interacted with FXR1, FXR2 and FMRP (Fig. 2B) as previously shown (Huang *et al*, 2020a; Marmor-Kollet *et al*, 2020; Sanders *et al*, 2020) and with some FG-Nups (detected by the monoclonal antibody mAb414) that are known to contribute to the biogenesis of the NPC in yeast (Onischenko *et al*, 2017) (Fig. 2B). Since the mAb414 is known to interact primarily with Nup62, as well as with Nup358, Nup214, and Nup153 (Davis & Blobel, 1987), it appears that UBAP2L may preferentially interact with Nup214 (Fig. 2B). Finally, ectopically expressed GFP-FXR1 interacted with Y-complex Nups and with UBAP2L (Fig. 2C). Taken together, the interaction of UBAP2L with Y-complex Nups as well as with the nuclear and cytoplasmic NPC assembly factors suggests a possible function of UBAP2L on Nups assembly and/or on NPC biogenesis.

**Fig. 2.**
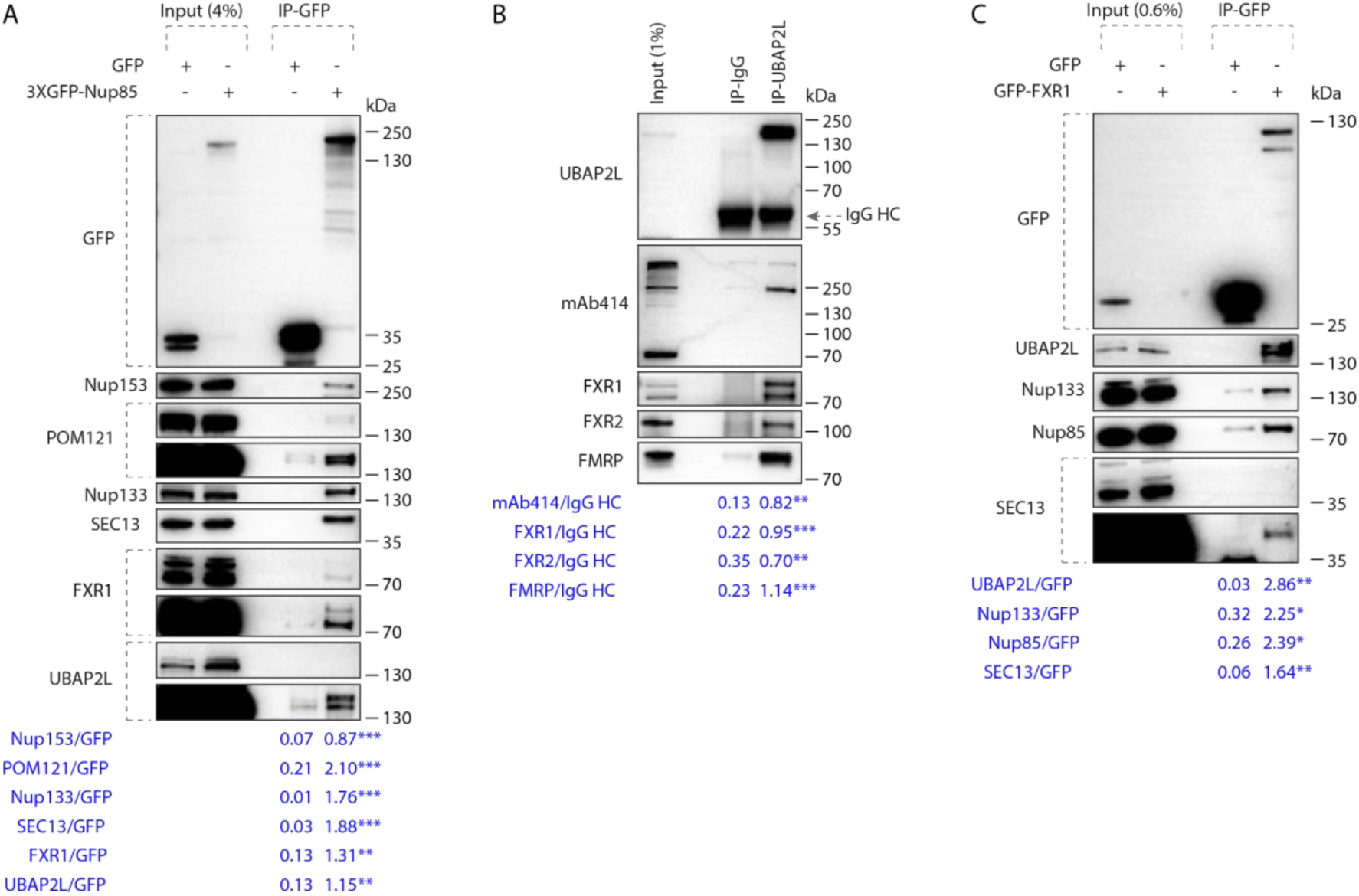
UBAP2L interacts with Nups and NPC assembly factors. **(A)** HeLa cells lysates expressing GFP alone or 3XGFP-Nup85 for 27h were immunoprecipitated using agarose GFP-Trap A beads (GFP-IP), analyzed by Western blot and signal intensities were quantified (shown a mean value, **P < 0.01, ***P < 0.001; *N* = 3). Molecular weight markers are indicated in kilodalton (kDa). **(B)** HeLa cells lysates were immunoprecipitation using UBAP2L antibody or IgG, analyzed by Western blot and signal intensities were quantified (shown a mean value, **P < 0.01, ***P < 0.001; *N* = 3). The arrow indicates the band corresponding to the IgG heavy chain (HC). **(C)** Lysates of HeLa cells expressing GFP alone or GFP-FXR1 for 27h were immunoprecipitated using agarose GFP-Trap A beads (GFP-IP), analyzed by Western blot and signal intensities were quantified (shown a mean value, *P < 0.05, **P < 0.01; *N* = 3).

### UBAP2L regulates Nups localization

To understand if UBAP2L regulates Nups assembly, we generated two clonal HeLa cell lines with CRISPR/Cas9-mediated deletion of the *UBAP2L* gene which were recently characterized (Guerber *et al*, 2023). As expected (Cirillo *et al*, 2020; Huang *et al*, 2020a; Youn *et al*, 2018), deletion of UBAP2L inhibited formation of SGs upon stress (Fig. S1E) and abolished nuclear localization of endogenous UBAP2L (Fig. S2B), confirming the specificity of UBAP2L antibodies. Relative to isogenic control cell line (wild type, (WT)), both UBAP2L knock-out (KO) cell lines revealed accumulation of foci containing Nups (Nup133, FG-Nups and RanBP2) as well as Importin-β and Exportin-1 in the cytoplasm but did not show defects in the localization of the NPC basket component Nup153 (Fig. 3, A to E). UBAP2L KO cells also displayed cytoplasmic granules containing both Importin-β and Nup133 (Fig. 3A) and likewise, RanBP2-containing granules co-localized with FG-Nups labelled by mAb414 (Fig. 3A). Such accumulation of cytoplasmic Nups strongly resembles the cellular phenotypes observed upon downregulation of the factors required for the assembly of NPCs at the NE such as FXR1 (Agote-Aran *et al*, 2020). We were unable to detect any changes in protein levels of several Nups as well as in Exportin-1 and Lamin A and B1 (Fig. 3F) in the whole cell extracts but deletion of UBAP2L led to reduced NE intensity of FG-Nups (Fig. 3, G and H). Fractionation experiments confirmed moderately reduced levels of Nups in the nucleus and an increased pool of cytoplasmic Nups upon deletion of UBAP2L (Fig. 3I), suggesting that UBAP2L does not regulate total protein levels of Nups but rather their localization to the NE during interphase. Owing to the fact that UBAP2L deletion can delay mitotic exit (Guerber *et al*, 2023), which could theoretically influence the length of G1 phase and, indirectly, the localization of Nups, we have arrested cells in G1 using lovastatin, which inhibits proteasome leading to the accumulation of p21 and p27 (Rao *et al*, 1999). Deletion of UBAP2L in G1-arrested cells led to accumulation of cytoplasmic Nup-containing granules, reduced NE intensity of FG-Nups (Fig. S2, C to F) without affecting the nuclear size (Fig. S2E). The same results were obtained in G0/G1-arrested cells using Psoralidin, which was suggested to transcriptionaly regulate cdk inhibitors (Gulappa *et al*, 2013) (Fig. S2, G to J). Lovastatin led to a decrease in nuclear size (Fig. S2F) as previously demonstrated (Iida *et al*, 2022) relative to Psoralidin (Fig. S2J) and to untreated HeLa cells (Fig. S5E) (Guerber *et al*, 2023) but no significant differences could be detected between WT and UBAP2L KO cells upon both treatments and under untreated conditions (Fig. S2, F and J, and Fig. S5E), which is in accordance with our recent published findings (Guerber *et al*, 2023). These results suggest that UBAP2L may regulate Nups without affecting the size of the nucleus and possibly the length of G1 phase. Our results demonstrate that UBAP2L localizes to the NE and the NPCs, interacts with Nups and regulates their localization and it may be involved in the assembly of the cytoplasmic Nups at the NE during interphase.

**Fig. 3.**
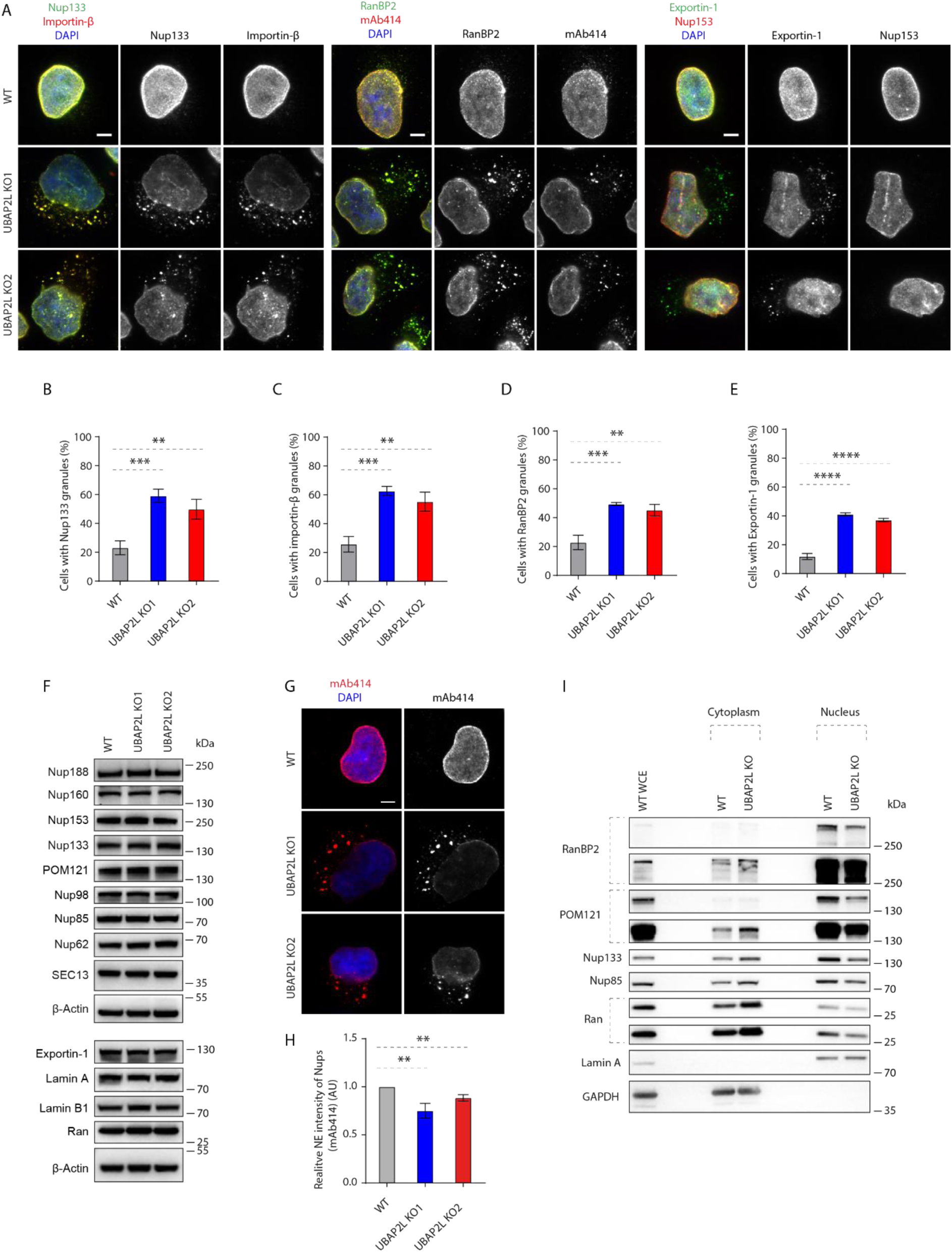
UBAP2L regulates Nups localization. (**A to E**) Representative immunofluorescence images depicting the localization of Nups and NPC-associated factors in wild type (WT) and UBAP2L Knock-out (KO) HeLa cells synchronized in interphase by double thymidine block and release (DTBR) at 12h (**A**). Nuclei were stained with DAPI. The percentage of cells with the cytoplasmic granules containing Nup133 (**B**), Importin-β (**C**), RanBP2 (**D**) and Exportin-1 (**E**) in (**A**) were quantified. At least 200 cells per condition were analyzed (mean ± SD, **P < 0.01, ***P < 0.001, ****P < 0.0001, two-tailed t-test, *N* = 3). Scale bars, 5 μm. **(F)** The protein levels of Nups and NPC-associated factors in WT and UBAP2L KO HeLa cells synchronized in interphase by DTBR at 12h were analyzed by Western blot. (**G and H**) Representative immunofluorescence images of FG-Nups (mAb414) at the NE in WT and UBAP2L KO HeLa cells in interphase cells synchronized by DTBR at 12h (**G**). Nuclei were stained with DAPI. The NE intensity of Nups (mAb414) in (**G**) was quantified (**H**). At least 150 cells per condition were analyzed (mean ± SD, **P < 0.01, two-tailed t-test, *N* = 3). Scale bar, 5 μm. **(I)** The nuclear and cytoplasmic protein levels of Nups and NPC transport-associated factors in WT and UBAP2L KO HeLa cells synchronized in the G1/S transition phase by thymidine 18h were analyzed by Western blot. WCE indicates whole cell extract.

### UBAP2L regulates localization of Nups in interphase but not in postmitotic cells

Two distinct pathways of NPC assembly at the NE have been described during the cell cycle in higher eukaryotic cells (Weberruss & Antonin, 2016). In the postmitotic pathway, NPC assembly occurs on segregated chromosomes, while during interphase, both Nup153 and POM121 drive *de novo* assembly of NPCs into an enclosed NE (D’Angelo *et al*, 2006; Doucet *et al*, 2010; Vollmer *et al*, 2015), which can be facilitated by FXR1 and microtubule-dependent transport of cytoplasmic Nups towards NE (Agote-Aran *et al*, 2020; Agote-Arán *et al*, 2021; Holzer & Antonin, 2020). Given the strong interaction of UBAP2L with FXR1 (Fig. 2, B and C), we hypothesized that UBAP2L may selectively affect Nups assembly during interphase. Indeed, accumulation of Nup-containing cytoplasmic granules could be first observed during late telophase, early G1 as well as in phospho-Rb-positive cells (which is present in mid-late G1, S and G2 phases) and but not during anaphase and early telophase stages (Fig. 4, A to F). FG-Nups assembled normally on segregating chromosomes in anaphase and on decondensing chromatin during early telophase (Fig. 4G) upon deletion of UBAP2L but reduced NE levels of FG-Nups were observed in early G1 and in phospho-Rb-positive cells in the absence of UBAP2L (Fig. 4, H and I). The percentage of cells in mid-late G1, S and G2 phases was not affected by UBAP2L deletion (Fig. 4J), further suggesting that the progression through interphase occurred normally in UBAP2L KO cells. Our findings suggest that UBAP2L drives Nups localization to NE during interphase but not in postmitotic cells.

**Fig. 4.**
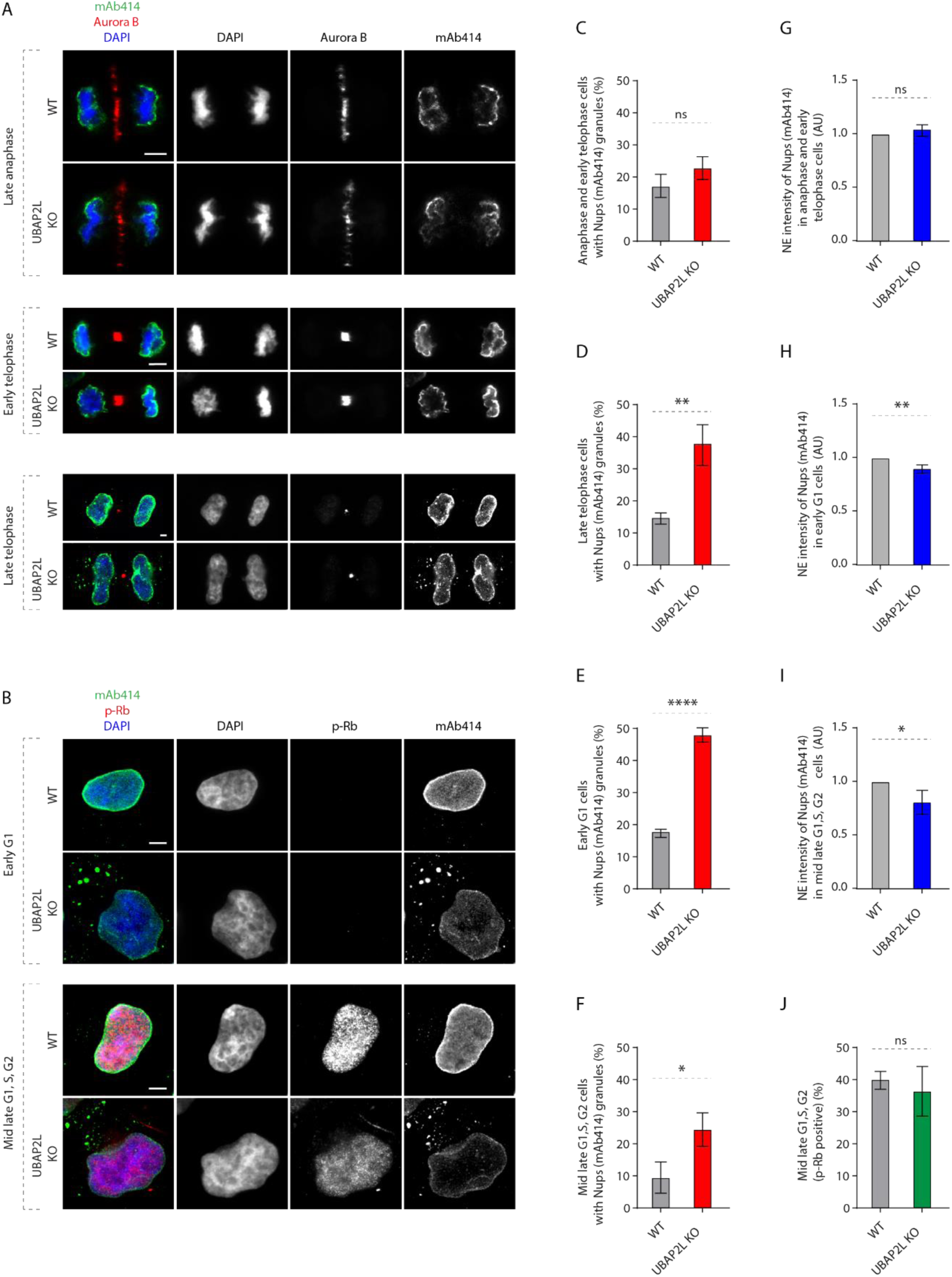
UBAP2L regulates localization of Nups in interphase but not in postmitotic cells. (**A and B**) Representative immunofluorescence images depicting the localization of Nups (mAb414) in WT and UBAP2L KO HeLa cells in different cell cycle stages. Mitotic cells were labeled by Aurora B (**A**) while p-Rb was used to distinguish between early G1 (p-Rb-negative cells) and mid-late G1, S and G2 (p-Rb-positive cells) stages (**B**). Nuclei were stained with DAPI. Scale bars, 5 μm. (**C to F**) The percentage of cells with the cytoplasmic granules of Nups (mAb414) in anaphase and early telophase (**C**), late telophase (**D**), early G1 (**E**) and mid-late G1, S, G2 (**F**) in (**A, B**) were quantified. At least 150 cells per condition were analyzed (mean ± SD, ns, non-significant, *P < 0.05, **P < 0.01, ****P < 0.0001, two-tailed t-test, *N* = 3). (**G to I**) The NE intensity of Nups (mAb414) in anaphase and early telophase cells (**G**), early G1 cells (**H**) and mid-late G1, S, G2 cells (**I**) in (**A, B**) were quantified. At least 100 cells per condition were analyzed (mean ± SD, ns, non-significant, *P < 0.05, **P < 0.01, two-tailed t-test, *N* = 3). **(J)** The percentage of p-Rb-positive cells in (**B**) was quantified. At least 150 cells per condition were analyzed (mean ± SD, ns, non-significant, two-tailed t-test, *N* = 3).

### UBAP2L mediates the assembly of the NPC scaffold elements and the biogenesis of NPCs

Our data demonstrate that UBAP2L deletion leads to decreased Nup levels at the NE and to the formation of Nup-containing granules in the cytoplasm. However, can UBAP2L also regulate the assembly of functional NPCs at the NE? The splitSMLM analysis revealed that deletion of UBAP2L decreased the density of the NPCs at the NE (Fig. 5, A and B) and confirmed the presence of RanBP2 and FG-Nups cytoplasmic assemblies (Fig. S3A), which often displayed linear-like organization with symmetrical RanBP2 distribution (Fig. S3A), contrary to the non-symmetrical distribution at the cytoplasmic site of the NE (Fig. 5A). Overexpression of Flag-tagged version of UBAP2L in interphase HeLa cells was also sufficient to moderately increase the density of NPCs at the NE (Fig. S3B and Fig. 5C), suggesting that UBAP2L might be required for NPC biogenesis onto intact NE. Flag-UBAP2L also occasionally co-localized with the cytoplasmic FG-Nups assemblies (Fig. S3B). The alignment and segmentation analysis of Nup133 or RanBP2 particles was performed as described previously (Andronov *et al*, 2022) and further suggested that the structure of the NE-localized NPCs was slightly altered upon deletion of UBAP2L (Fig. 5, A and D). Relative to WT, UBAP2L KO cells showed moderately increased percentage of NPCs with a 4-fold rotational symmetrical arrangement of the scaffold spokes, while the number of NPC structures with 5 to 8-fold symmetrical organization was slightly decreased upon UBAP2L deletion (Fig. 5, A and D). Two clonal U2OS cell lines with CRISPR/Cas9-mediated deletion of *UBAP2L* gene with stably integrated Nup96-GFP (Nup96-GFP knock-in (KI)) (Fig. S4, A and B) likewise showed the accumulation of cytoplasmic Nup-containing granules (Nup96-GFP and FG-Nups) (Fig. S4, C to E) and reduced density of the NPCs at the NE (Fig. 5, E and F).

**Fig. 5.**
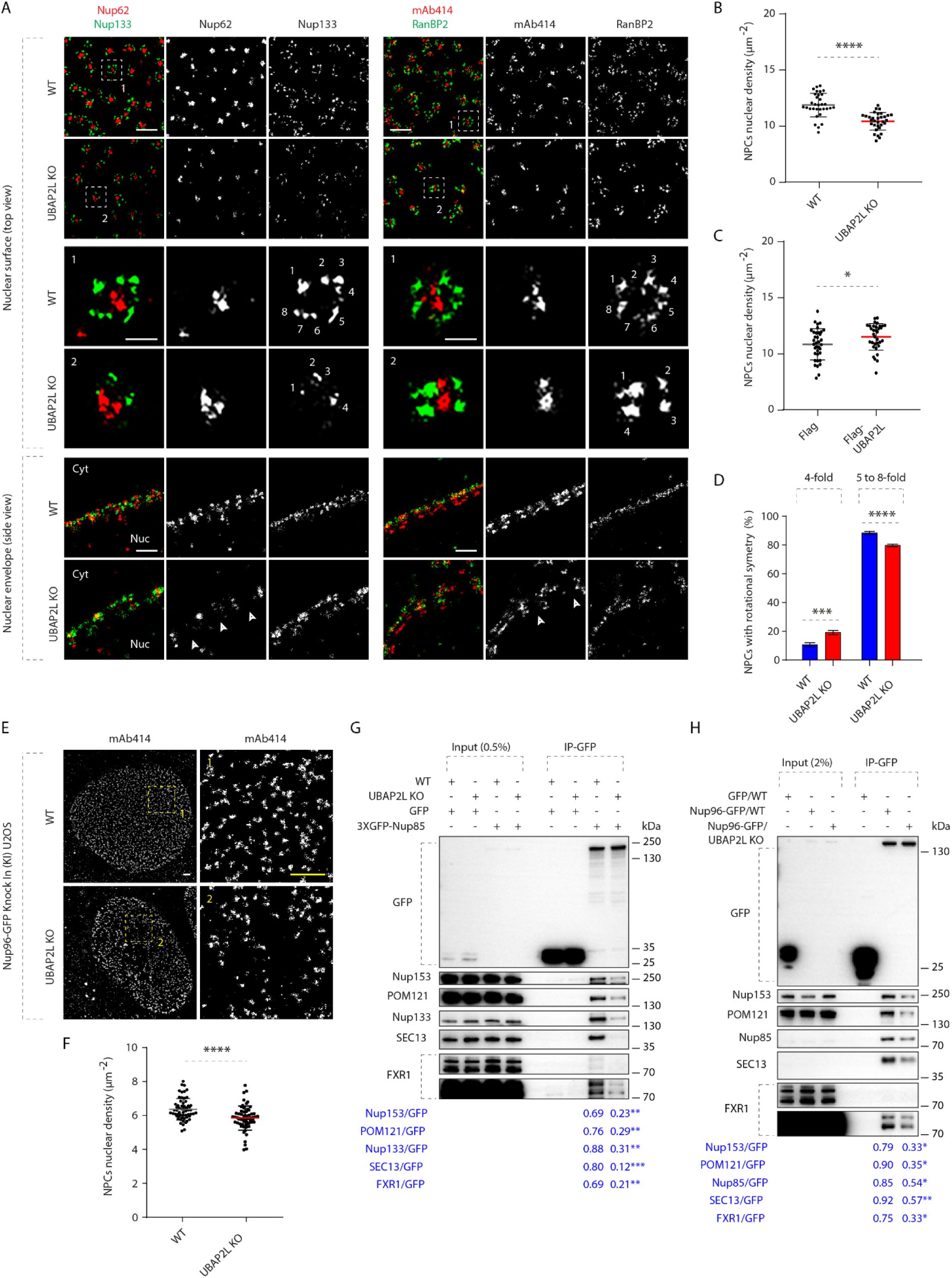
UBAP2L mediates the assembly of the NPC scaffold elements and the biogenesis of NPCs. (**A**) Representative splitSMLM images depicting several NPC components on the nuclear surface (top view) and in the cross section of the NE (side view) in WT and UBAP2L KO HeLa cells synchronized in early interphase by DTBR at 12h. Nup133 signal labels the cytoplasmic and nuclear rings of the NPC, the localization of the central channel is visualized by Nup62 and mAb414 antibodies and cytoplasmic filaments are labeled by RanBP2. The magnified framed regions are shown in the corresponding numbered panels. Nuclear (Nuc) and cytoplasmic (Cyt) side of the NE are indicated in the side view. The arrowheads indicate the disrupted localization of Nup62 or mAb414 at NE in UBAP2L KO HeLa cells and the numbers point to the individual identified spokes of the NPC. Scale bars, 300 and 100 nm, respectively. (**B and C**) The nuclear density of NPCs (mAb414 and RanBP2) in cells shown in (A) was quantified (**B**) (mean ± SD, ****P < 0.0001, two-tailed t-test; counted 32 cells per cell line). The nuclear density of NPCs (mAb414) in HeLa cells expressing Flag alone or Flag-UBAP2L for 35h and synchronized in interphase by DTBR at 12h was quantified (**C**) (mean ± SD, *P < 0.05, two-tailed t-test; counted 36 cells for Flag and 33 cells for Flag-UBAP2L). The corresponding representative images are shown in the Fig. S3B. (**D**) The 8-fold rotational symmetry of NPCs in (**A**) was quantified by alignment of Nup133 particles and segmentation analysis (mean ± SD, ***P < 0.001, ****P < 0.0001, two-tailed t-test; counted 851 NPCs for WT HeLa cell line and 559 NPCs for UBAP2L KO HeLa cell line). (**E and F**) Representative SMLM immunofluorescence images of FG-Nups (mAb414) at the nuclear surface in Nup96-GFP KI U2OS WT and UBAP2L KO cells in interphase cells synchronized by DTBR at 12h (**E**). The nuclear density of NPCs (mAb414) in cells shown in (**E**) was quantified in (**F**) (mean ± SD, ****P < 0.0001, two-tailed t-test; counted 60 cells per cell line). Scale bars, 1 μm. (**G and H**) Lysates of interphase WT and UBAP2L KO HeLa cells expressing GFP alone or 3XGFP-Nup85 for 27h were immunoprecipitated using agarose GFP-Trap A beads (GFP-IP), analyzed by Western blot and signal intensities were quantified (shown a mean value, **P < 0.01, ***P < 0.001; *N* = 3) (**G**). Lysates of interphase U2OS cells expressing GFP alone for 27h and Nup96-GFP KI U2OS WT and UBAP2L KO cells were immunoprecipitated using agarose GFP-Trap A beads (GFP-IP), analyzed by Western blot and signal intensities were quantified (shown a mean value, *P < 0.05, **P < 0.01; *N* = 3) (**H**).

Consistent with the observed role of UBAP2L in the biogenesis of mature NPCs, deletion of UBAP2L in HeLa cells reduced the interaction of GFP-Nup85 with other components of the Y-complex, Nup133 and SEC13 in both unsynchronized (Fig. 5G) and in G1/S-synchronized cells (Fig. S4F) as well as decreased the interaction of GFP-Nup85 with the two Nups, Nup153 and POM121 (Fig. 5G), involved in the assembly of the NPCs at the enclosed NE through the interphase pathway (Funakoshi *et al*, 2011; Vollmer *et al*, 2015). Immunoprecipitated (IP) Nup96-GFP in U2OS cells also demonstrated reduced interaction of Y-complex components Nup85 and SEC13 and inhibition of Nup96-GFP binding to Nup153 and POM121 in the absence of UBAP2L (Fig. 5H). Interestingly, the interaction of endogenous Nup85 with other components of the Y-complex appeared moderately increased in G1/S cells relative to cells arrested in prometaphase using Eg5 inhibitor STLC (Fig. S4G), suggesting that Y-complex assembly may also take place during interphase. In addition, the ‘interaction of the cytoplasmic Nup transporter factor FXR1 with both GFP-Nup85 and Nup96-GFP was reduced in the absence of UBAP2L (Fig. 5, G and H) and UBAP2L deletion inhibited the binding of immunoprecipitated GFP-FXR1 with Nup85, SEC13 and with the components of the dynein complex dynactin p150^Glued^ and BICD2 (Fig. S4H) that work with FXR1 to transport Nups along microtubules towards NE during interphase (Agote-Aran *et al*, 2020). Collectively, these results demonstrate that UBAP2L is critically involved in the biogenesis of NPCs at the NE during interphase likely through the regulation of the assembly of the NPC scaffold elements from the cytoplasmic Nups and by facilitating the interaction of the Y-complex with both the nuclear (Nup153, POM121) as well as with the cytoplasmic (FXR1, dynein complex) NPC assembly signals.

### UBAP2L regulates localization of the Nup transporting factor FXR1

What is the molecular, UBAP2L-dependent mechanism fueling the assembly of cytoplasmic Nups into Y-complex? The cellular phenotypes on cytoplasmic Nups observed upon deletion of UBAP2L strongly resemble downregulation of the fragile X-related proteins (FXRPs) (FXR1, FXR2 and FMRP) which drive transport and spatial assembly of the cytoplasmic Nups to the NE in human cells during early interphase (Agote-Aran *et al*, 2020; Agote-Arán *et al*, 2021). The fact that UBAP2L not only facilitated the interaction of the Y-complex with Nup153 and POM121 but also with FXR1 and the dynein complex (Fig. 5, G and H, and Fig. S4H) and that FXRPs strongly interacted with UBAP2L (Fig. 2, B and C) prompted us to analyze the dynamics of FXRPs in UBAP2L-deficient cells in more detail. Interestingly, deletion of UBAP2L led to changes in the localization of FXR1 protein. In contrast to WT cells where FXR1 was localized at the NE and diffusely in the cytoplasm, as reported previously (Agote-Aran *et al*, 2020), both UBAP2L KO cell lines displayed reduced NE localization of FXR1 and formation of cytoplasmic FXR1-containing granules (Fig. S5, A and C) in addition to FG-Nups-containing granules, which did not co-localize with FXR1 in the cytoplasm (Fig. S5, A and B). Both UBAP2L KO cell lines also showed irregular nuclear shape (Fig. S5D) but no changes in the nuclear size (Fig. S5E) could be observed, in accordance with our previous findings (Guerber *et al*, 2023). Fractionation experiments confirmed moderately reduced levels of FXR1 in the nucleus upon deletion of UBAP2L both in G1-synchronized (Fig. S5F) and in unsynchronized interphase cells (Fig. S5G), similar to Nups and to the nuclear transport factor Ran (Fig. S5, F and G). The same phenotype was observed for FMRP (Fig. S6, A and B), and UBAP2L deletion did not appear to affect protein levels of any of the three FXRPs (Fig. S6C). Downregulation of endogenous UBAP2L using specific siRNAs confirmed the cellular phenotypes of UBAP2L KO cells and displayed accumulation of FXR1 foci, cytoplasmic Nups-containing granules and irregular nuclear shape as also observed upon depletion of FXR1 and in contrast to control cells (Fig. S6, D to G). These results suggest that FXR1 cytoplasmic granules are not the result of any possible compensation effects due to stable deletion of UBAP2L in KO cells. Since UBAP2L was previously demonstrated to contribute to the assembly of SGs (Cirillo *et al*, 2020; Huang *et al*, 2020a; Youn *et al*, 2018) and FXRPs and Nups are able to localize to these protein assemblies (Huang *et al*, 2020a; Zhang *et al*, 2018), we aimed to understand if observed phenotypes could be linked to cellular stress signaling. As expected (Cirillo *et al*, 2020; Huang *et al*, 2020a; Youn *et al*, 2018), deletion of UBAP2L inhibited formation of SGs (Fig. S1E) upon stress but the SG components G3BP1 and TIA-1 did not localize to FXR1-containing granules under normal growing conditions in UBAP2L KO cells (Fig. S6, H and I), suggesting that FXR1 foci are distinct from SGs. Our findings indicate that UBAP2L-mediated regulation of Nups might be independent of UBAP2L’s function on SGs. Importantly, UBAP2L not only facilitates the interaction of FXRPs with the scaffold Nups but also helps to localize FXRPs to the NE thereby fueling the assembly of Nups from the cytoplasm to the nucleus.

### Arginines within the RGG domain of UBAP2L mediate the function of UBAP2L on Nups and FXRPs

To dissect the molecular basis of the UBAP2L-FXR1-Nup pathway and to understand if the function of UBAP2L on cytoplasmic Nups and on FXRPs is specific, we performed rescue experiments. In contrast to GFP, ectopic expression of GFP-UBAP2L efficiently rescued Nup-and FXR1-granules as well as the irregular nuclei phenotypes in both UBAP2L KO cell lines (Fig. S7, A to E). GFP-UBAP2L protein fragment encompassing 98-430 aa was required (Fig. S8, A and B) and sufficient (Fig. S8C) for the interaction with FXR1 in the immunoprecipitation experiments. Interestingly, the 98-430 aa protein fragment of UBAP2L contains the RGG domain (Fig. S8A) which often engages in interactions with mRNAs and mediates UBAP2L’s function in protein translation and RNA stability (Luo *et al*, 2020). Surprisingly, GFP-tagged UBAP2L (Fig. S8D) and endogenous UBAP2L (Fig. S8E) interacted with endogenous FXR1 and FMRP despite the absence of RNAs after RNase A treatment, suggesting that the role of UBAP2L on FXRPs-Nups pathway may be, to a large extent, RNA-independent. The arginines present in the RGG domains were previously demonstrated to regulate localization of other proteins also in an RNA-independent manner (Thandapani *et al*, 2013) and to be asymmetrically di-methylated (ADMA) by the protein arginine methyltranferase PRMT1 (Huang *et al*, 2020a; Maeda *et al*, 2016). Indeed, Flag-tagged mutant form of UBAP2L, where all 19 arginines were exchanged to alanines (UBAP2L R131-190A), did not interact with endogenous PRMT1 and showed reduced ADMA signal as expected (Fig. 6A). The UBAP2L R131-190A mutant also did not bind to Nups and FXR1 (Fig. 6A), suggesting the role of arginines within the RGG domain of UBAP2L in Nups assembly. The GFP-UBAP2L protein fragment encompassing 98-430 aa could rescue localization defects of Nups and FXR1 in UBAP2L KO cells, in a manner similar to the full length UBAP2L protein (Fig. S9, A to D) but the UBAP2L R131-190A mutant was unable to restore the FXR1 and Nups localization defects and irregular nuclear shape in UBAP2L KO cells (Fig. 6, B to E). We conclude that the function of UBAP2L on the regulation of FXRPs and Nups localization may be mediated through the arginines present within its RGG domain.

**Fig. 6.**
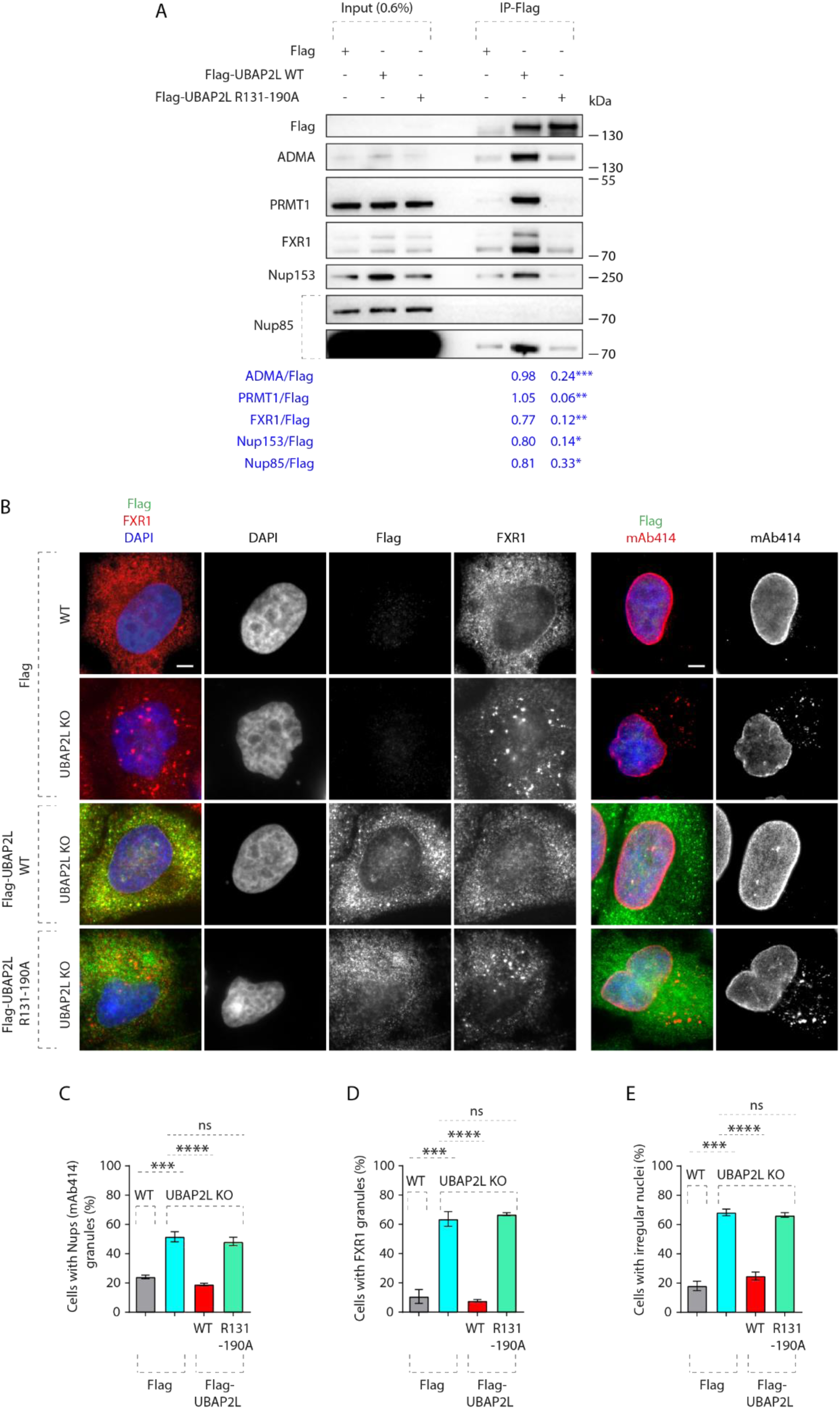
Arginines within the RGG domain of UBAP2L mediate the function of UBAP2L on Nups and FXRPs. (**A**) Lysates of interphase HeLa cells expressing Flag alone, Flag-UBAP2L WT or mutated Flag-UBAP2L version where 19 arginines located in the RGG domain were replaced by alanines (R131-190A) for 27h were immunoprecipitated using Flag beads (Flag-IP), analyzed by Western blot and signal intensities were quantified (shown a mean value, *P < 0.05, **P < 0.01, ***P < 0.001; *N* = *3*). (**B to E**) Representative immunofluorescence images depicting nuclear shape and localization of FXR1 and Nups (mAb414) in WT and UBAP2L KO HeLa cells expressing Flag alone or Flag-UBAP2L (WT or R131-190A) for 60h and synchronized in interphase by DTBR at 12h (**B**). Nuclei were stained with DAPI. The percentage of cells with the cytoplasmic granules of Nups (mAb414) (**C**) and of FXR1 (**D**) and irregular nuclei (**E**) shown in (**B**) were quantified. At least 200 cells per condition were analyzed (mean ± SD, ns: not significant, ***P < 0.001, ****P < 0.0001, two-tailed t-test, *N* = 3). Scale bars, 5 μm.

### UBAP2L regulates localization of FXR1 to the NE

How can the function of UBAP2L in NPC biogenesis and in ensuring the interaction of FXR1 with Nups be linked to the observed subcellular localization of FXR1? And why, and when can FXRPs form cytoplasmic assemblies in the absence of UBAP2L? Although UBAP2L regulates some factors involved in mitotic exit (Guerber *et al*, 2023; Maeda *et al*, 2016), the localization defects of Nups upon UBAP2L deletion could be also observed in cells arrested in G1 phase (Fig. S2, C to J). In addition, inhibition of Polo-like kinase 1 (PLK1) activity, the downstream target of UBAP2L during mitosis, was reported to rescue the mitotic defects observed in the absence of UBAP2L (Guerber *et al*, 2023) but it could not reverse the Nup localization defects in the same experimental setting (Fig. S10, A and B), arguing that UBAP2L-dependent regulation of Nups assembly could be largely uncoupled from the role of UBAP2L in mitotic progression.

Importantly, the increased numbers of FXR1-containing foci were also observed in UBAP2L KO late telophase cells when compared to the corresponding WT cells synchronized in the same cell cycle stage (Fig. S10, C and D). The average size of the FXR1-containing granules was likewise increased in late telophase synchronized UBAP2L KO relative to WT cells (0.346 and 0.218 μm^2^, respectively) (Fig. S10, C and E). Reduced NE localization of FXR1 and formation of cytoplasmic granules were observed in early and mid-late G1, S and in G2 phases in UBAP2L KO relative to WT cells (Fig. S10, F to I). In addition, endogenous UBAP2L could interact with endogenous FXR1 and FMRP in asynchronous cells as well as in cells synchronized during mitosis and in interphase (Fig. S11A). Interestingly, the effect of UBAP2L deletion on the percentage of FXR1 granules-containing cells, the number of granules per cell and the size of FXR1 granules was the most evident in early G1 compared to other cell cycle stages analyzed (Fig. S10, F to I), in line with our findings suggesting that UBAP2L preferentially regulates Nups localization to NE during early G1 (Fig. 4, A to I). The fact that FXR1-containing granules are also observed in the WT late telophase cells, although to a lesser extent as compared to UBAP2L KO cells (Fig. S10, C to E), suggests that these structures do not form *de novo* upon deletion of UBAP2L but may originate from some similar assemblies existing before mitotic exit.

For this reason, we analyzed FXR1 and FMRP localization during mitosis in wild-type cells synchronized in prometaphase-like stage using Nocodazole or Eg5 inhibitor STLC. Interestingly, wild-type mitotic cells displayed strong accumulation of granules containing both FXR1 and FMRP (Fig. 7A). Time-lapse analysis using live video spinning disk microscopy of cells expressing GFP-FXR1 revealed its dynamic localization during mitotic progression and confirmed the presence of GFP-FXR1-containing granules in control mitotic cells (Fig. 7, B to D) where GFP-FXR1 granules could be observed first during late prophase and throughout prometaphase, metaphase and anaphase stages. Interestingly, unlike in control cells where GFP-FXR1 mitotic granules spread out in the vicinity of the NE concomitant with the nuclei reformation during mitotic exit, in UBAP2L-deleted cells, these granules remained in the cytoplasm, surrounding the nucleus and GFP-FXR1 localization at the NE appeared to be reduced (Fig. 7, B to D). Accordingly, both the number as well as the average size of FXR1-containing granules were increased in dividing UBAP2L-deficient cells relative to WT cells (Fig. 7, C and D). These results suggest that UBAP2L may remodel FXR1 protein assemblies present in mitotic cells to restrict and ensure their timely localization to the vicinity of the NE after completion of mitosis, where it could interact with Nups and transport them towards NE allowing for the formation of mature NPCs during early interphase. Indeed, endogenous UBAP2L and FXR1 can localize to NE and occasionally co-localize in the cytoplasmic assemblies in the proximity of NE in early interphasic cells (Fig. 7E). In addition, Flag-tagged WT, but not the R131-190A mutant form of UBAP2L, frequently co-localized to FXR1-containing granules in the proximity of NE in late telophase cells (Fig. 7F) and WT but not R131-190A mutant UBAP2L was able to disperse endogenous FXR1-containing mitotic granules (Fig. S11, B to D). Similar observations were made when either the full length or the 98-430 aa UBAP2L-fragment fused to GFP, but not GFP alone, were expressed in STLC-synchronized mitotic cells (Fig. S11, E to G), suggesting that UBAP2L may chaperone and/or remodel FXR1 to ensure its interaction with Nups and their timely localization to the NE. The exact molecular mechanism underlying UBAP2L-mediated remodeling of FXR1 will have to be investigated in the future but it is interesting to note that DNAJB6, a molecular chaperone of the heat shock protein network, which was demonstrated to prevent aggregation of Nups and promote their NE assembly during interphase (Kuiper *et al*, 2022) could also interact with endogenous UBAPL2 in our hands (Fig. S11H), further corroborating the role of UBAP2L in the assembly of cytoplasmic Nups. Collectively, our results identify UBAP2L as an important component of the FXRPs-Nups pathway that drives assembly of NPCs during early interphase by regulating the localization of FXR1 and Nups to the NE during early G1.

**Fig. 7.**
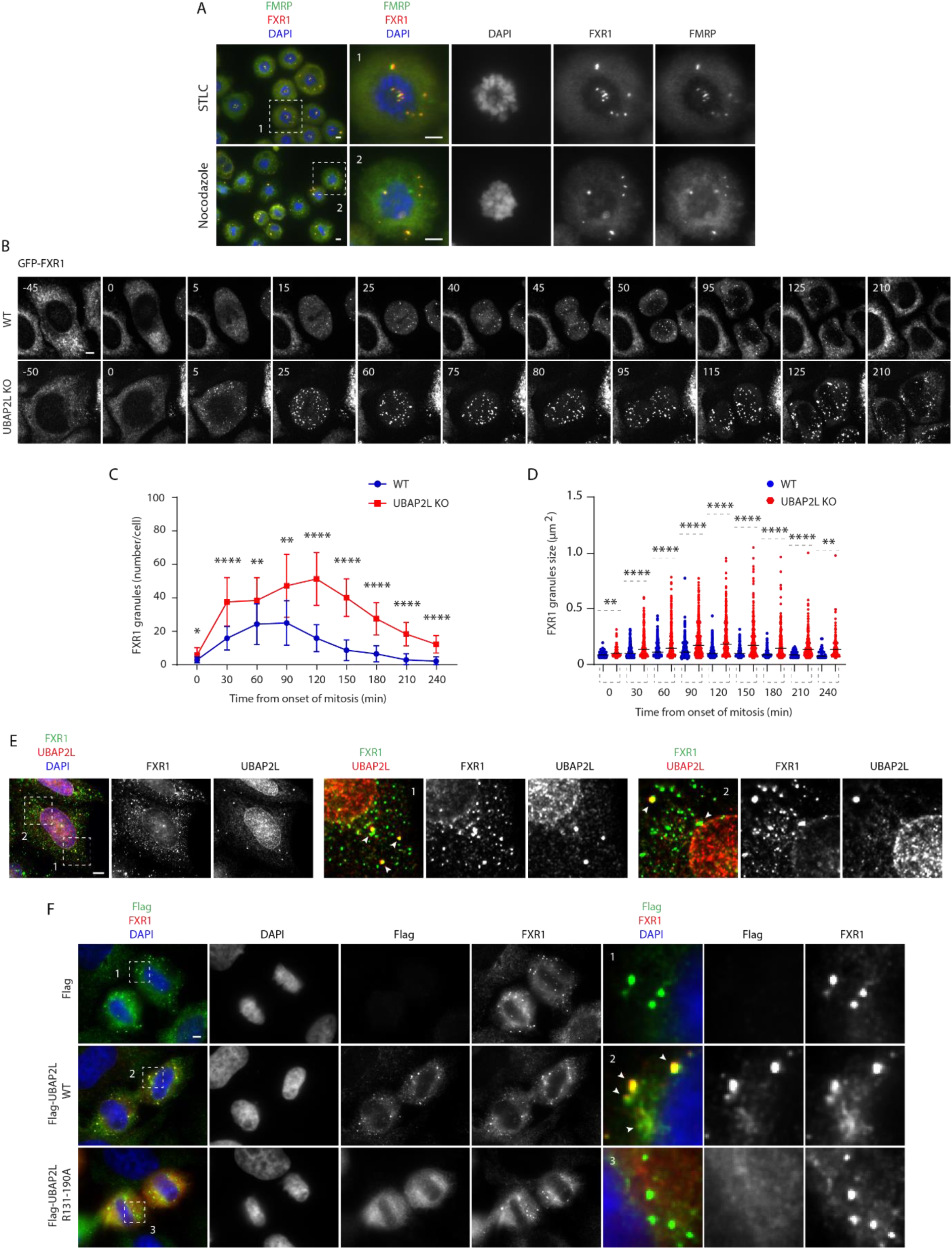
UBAP2L remodels FXR1-protein assemblies in the cytoplasm and drives localization of FXR1 to the NE. (**A**) Representative immunofluorescence images depicting the localization of FXR1 and FMRP in HeLa cells synchronized in prometaphase using STCL 16h or nocodazole 16h. Chromosomes were stained with DAPI. Scale bars, 5 μm. (**B to D**) WT and UBAP2L KO HeLa cells expressing GFP-FXR1 were synchronized by DTBR and analyzed by live video spinning disk confocal microscopy. The selected representative frames of the movies are depicted and time is shown in minutes. Time point 0 indicates mitotic entry during prophase (**B**). GFP-FXR1 granules number (number/cell) shown in (**B**) at indicated times during mitotic progression were quantified (**C**). GFP-FXR1 granules sizes (granule ≥ 0.061 µm^2^) shown in (**B**) at indicated times during mitotic progression were quantified (**D**). 16 WT and 11 UBAP2L KO HeLa cells were counted, respectively. Scale bar, 5 μm. (**E**) Representative immunofluorescence images depicting the cytoplasmic and NE localization of endogenous UBAP2L and FXR1 in interphase HeLa cells. Nuclei were stained with DAPI. The magnified framed regions are shown in the corresponding numbered panels. The arrows indicate co-localization of UBAP2L and FXR1 foci in the cytoplasm. Scale bar, 5 μm. (**F**) Representative immunofluorescence images depicting the localization of FXR1, Flag alone and Flag-UBAP2L (WT or R131-190A) in late telophase in HeLa cells. Nuclei were stained with DAPI. The magnified framed regions are shown in the corresponding numbered panels. Note that Flag-UBAP2L WT but not Flag alone and Flag-UBAP2L R131-190A, is localized to FXR1 containing granules in proximity of NE. Scale bar, 5 μm.

### UBAP2L-mediated biogenesis of NPCs ensures nuclear transport, adaptation to nutrient stress and cellular proliferation

Next, it was important to understand the physiological relevance and functional implications of the UBAP2L-mediated assembly of NPCs at the NE. One of the main functions of the NPCs is the regulation of the nucleocytoplasmic transport across the NE. Our data so far demonstrated that deletion of UBAP2L leads to the cytoplasmic sequestration of FG-Nups (Fig. 3, A and G), which constitute the selective permeability barrier of NPCs as well as of Importin-β and Exportin-1 (Fig. 3, A and C, and E), the essential components of the nucleocytoplasmic transport system (Pemberton & Paschal, 2005). UBAP2L KO cells also display a reduced number of NPCs at the intact NE (Fig. 5, A and B, E and F). To understand if these Nups and NPCs defects in UBAP2L-deficient cells affect the function of nuclear pores, we measured the rates of nucleocytoplasmic transport of an ectopic import/export reporter plasmid XRGG-GFP that shuttles to the nucleus (accumulating in the nucleoli) when induced with dexamethasone as previously described (Agote-Aran *et al*, 2020; Love *et al*, 1998). Deletion of UBAP2L decreased the rates of XRGG-GFP nuclear import (Fig. 8, A and B) and its nuclear export (Fig. 8, C and D) relative to WT cells, suggesting that UBAP2L is important for the transport function of NPCs. To corroborate these observations using a marker which does not localize at specific structures, we analyzed the gradient of endogenous Ran, a guanine nucleotide triphosphatase, as shown in previously (Coyne *et al*, 2020; Zhang *et al*, 2015). Most of Ran protein is actively imported to the nucleus with help of transport factors, a process that requires Ran binding to GDP (Ribbeck *et al*, 1998; Smith *et al*, 2002, 1998). Therefore, we analyzed the nuclear-cytoplasmic (N/C) distribution of Ran and observed that UBAP2L deletion led to significant reduction in the N/C ratio of Ran (Fig. 8, E and F). Together, with our analysis in living cells, and with the reduced nuclear levels of Ran in fractionation experiments (Fig. 3I, and Fig. S5, F and G), these results suggest that UBAP2L may facilitate the nucleocytoplasmic transport across the NE.

**Fig. 8.**
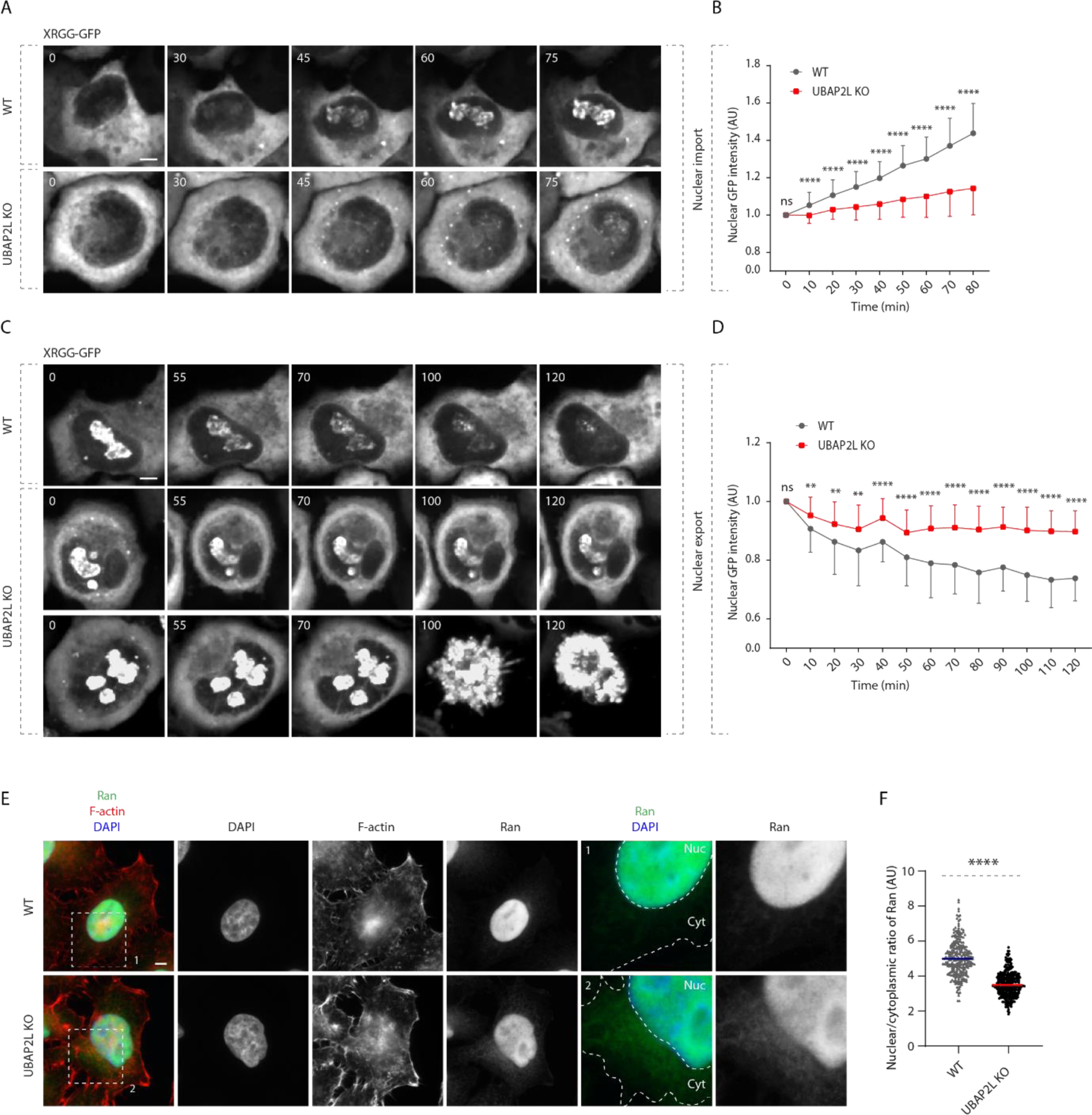
UBAP2L regulates nucleocytoplasmic transport. (**A to D**) WT and UBAP2L KO HeLa cells expressing reporter plasmid XRGG-GFP for 30h were analyzed by live video spinning disk confocal microscopy. The selected representative frames of the movies are depicted and time is shown in minutes. Time point 0 in top panel (nuclear import of XRGG-GPF) indicates that dexamethasone (0.01 μM) was added, while time point 0 in bottom panel (nuclear export of XRGG-GPF) indicates that dexamethasone was washed out (**A, B**). The arrowheads indicate dead cells in UBAP2L KO cells. The nuclear intensity (fold change) of XRGG-GFP (to DNA labeled by SiR-DNA probe) in top panel (nuclear import) (**C**) and in bottom panel (nuclear export) (**D**) shown in (**A, B**) were quantified. At least 10 cells per condition were analyzed (mean ± SD, ns: not significant, **P < 0.01, ****P < 0.0001, two-tailed t-test, *N* = 3). Scale bars, 5 μm. (**E and F**) Representative immunofluorescence images depicting the nuclear (Nuc) and cytoplasmic (Cyt) localization of Ran in asynchronously proliferating WT and UBAP2L KO HeLa cells (**E**). Nuclei were stained with DAPI. Actin filaments (also known as F-actin) were stained with phalloidin. The magnified framed regions are shown in the corresponding numbered panels. The nuclear (Nuc)-to-cytoplasmic (Cyt) ratio of Ran shown in (**E**) was quantified (**F**) (mean ± SD, ****P < 0.0001, two-tailed t-test; counted 277 cells for WT and 306 cells for UBAP2L KO). Scale bars, 5 μm.

Interestingly, in the live video analysis we observed that UBAP2L-deficient cells displaying strong transport defects may undergo cellular death (Fig. 8C) in accordance with the previous reports demonstrating an essential role of transport across NE for cell viability (Hamada *et al*, 2011). Colony formation assays showed that the long-term proliferation capacity of both UBAP2L KO cell lines was reduced relative to WT cells (Fig. S12, A to E) in agreement with our published study (Guerber *et al*, 2023) and propidium iodide labelling and flow cytometry indicated reduced viability of UBAP2L KO cells (Fig. S12, F and G). Future studies will have to address whether UBAP2L-dependent regulation of NPC assembly can directly promote cell survival or if the effects of UBAP2L deletion on NPC function and viability are circumstantial. Because the Y-complex can selectively affect survival and proliferation of cancer cells in response to presence of nutrients (such as high serum and growth factors) (Sakuma *et al*, 2020) and changes in nutrient availability can lead to NPC reorganization (clustering) in fission yeast (Varberg *et al*, 2021), we studied how UBAP2L-dependent biogenesis of NPCs can be affected by nutrient deprivation in human cells.

Nutrient deprivation further potentiated inhibition of cell viability in UBAP2L-dependent manner (Fig. S12, F and G) and led to reduced NE levels of Nups and accumulation of Nup-containing granules (Fig. 9, A to C). Interestingly, NE localization and protein levels of UBAP2L were moderately reduced upon nutrient deprivation (Fig. 9, A and D, and E) but the total protein levels of several tested Nups were unaffected under nutrient poor conditions (Fig. 9E).

**Fig. 9.**
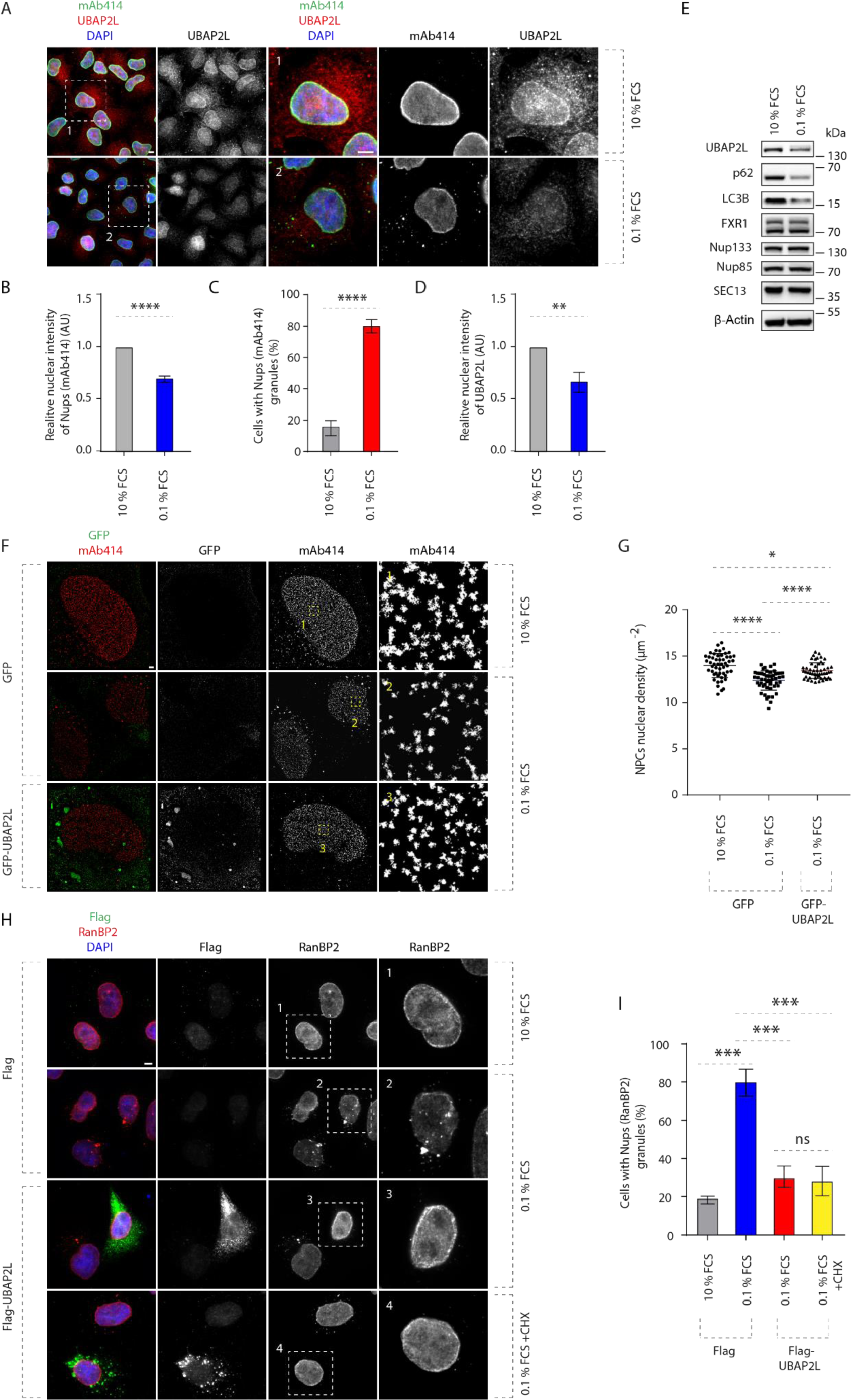
UBAP2L ensures NPC biogenesis upon nutrient stress. (**A to D**) Representative immunofluorescence images depicting the localization of UBAP2L and Nups (mAb414) in HeLa cells cultured in the indicated concentrations of serum for 72h (**A**). Nuclei were stained with DAPI. Scale bars, 5 μm. The nuclear intensity of Nups (mAb414) (**B**) the percentage of cells with the cytoplasmic granules of Nups (mAb414) (**C**) and nuclear intensity of UBAP2L (**D**) shown in (**A**) were quantified. At least 150 cells per condition were analyzed (mean ± SD, **P < 0.01, ***P < 0.001, ****P < 0.0001, two-tailed t-test, *N* = 3). (**E**) The protein levels of UBAP2L, Nups, FXR1 and other indicated factors in HeLa cells cultured in the indicated concentrations of serum for 72h were analyzed by Western blot. (**F and G**) Representative SMLM immunofluorescence images of FG-Nups (mAb414) at the nuclear surface in interphase HeLa cells expressing GFP alone or GFP-UBAP2L WT for 48h cultured in the indicated concentrations of serum for 72h (**F**). The magnified framed regions are shown in the corresponding numbered panels. The nuclear density of NPCs (mAb414) in cells shown in (**F**) was quantified (**G**) (mean ± SD, *P < 0.05, ****P < 0.0001, two-tailed t-test; counted 51 cells per cell line). Scale bar, 1 μm. (**H and I**) Representative immunofluorescence images depicting the localization of RanBP2 in HeLa cells expressing Flag alone or Flag-UBAP2L for 30h cultured in the indicated concentrations of serum for 72h (**H**). Note that Cycloheximide (CHX) was used at a concentration of 0.1 mg/ml for 8h prior to sample collection. The magnified framed regions are shown in the corresponding numbered panels. Nuclei were stained with DAPI. The percentage of cells with the cytoplasmic granules containing RanBP2 shown in (**H**) was quantified (**I**). At least 200 cells per condition were analyzed (mean ± SD, ns: not significant, ***P < 0.001, two-tailed t-test, *N* = 3). Scale bar, 5 μm.

Nutrient stress could also lead to reduced density of NPCs at the NE, a phenotype which could be partially rescued by overexpression of GFP-UBAP2L (Fig. 9, F and G), suggesting that presence of UBAP2L is important for NPC biogenesis also under nutrient stress conditions. Finally, nutrient deprivation could induce the formation of the cytoplasmic Nup granules, which were rescued by Flag-UBAP2L overexpression also upon inhibition of active protein translation (using cycloheximide) (Fig. 9, H and I), suggesting that UBAP2L-mediated NPC formation under nutrient stress conditions is independent of production of new proteins at least during early interphase. The possible regulation of NPC biogenesis by UBAP2L in response to nutrient poor conditions or upon induction of autophagy will have to be investigated in future. Taken together, our data are consistent with the hypothesis that the role of UBAP2L in NPC biogenesis at the NE is important for nuclear transport and adaptation to nutrient stress.

## Discussion

NPCs are large (approximately 100 nm wide and 40 nm high) eightfold symmetrical assemblies composed of more than 550 copies of around 30 different Nups. Nups assemble into biochemically stable subcomplexes that form eight identical protomer units, traditionally named “spokes” which are radially arranged around the central channel or the “central transporter”. Although deviations from typical eightfold rotational symmetry have been observed in Xenopus oocytes (Hinshaw & Milligan, 2003) and NPCs can dilate their inner ring by moving the spokes away from each other (Mosalaganti *et al*, 2018), the identity of the molecular pathways defining the NPC structural organization remains unknown.

Our data suggest a model (Fig. 10) how UBAP2L ensures assembly of the NPC scaffold elements into mature NPCs at the intact NE in human cells during early G1. On one hand, UBAP2L localizes to the NE and to NPCs and drives the formation of the Y-complex and its interaction with Nup153 and POM121, which are known to be crucial for the Y-complex recruitment to the NE during interphase (Funakoshi *et al*, 2011; Vollmer *et al*, 2015). On the other hand, UBAP2L can remodel or “chaperone” FXRP proteins to restrict their timely localization to the NE and their interaction with the Y-complex. Thus, UBAP2L can link and integrate the nuclear and the cytoplasmic NPC upstream assembly signals during interphase. Since FXRPs were shown to transport cytoplasmic Y-complex Nups towards the NE through a microtubule-based mechanism (Agote-Aran *et al*, 2020), their interaction with UBAP2L may fuel the assembly of Y-complexes and biogenesis of new NPCs with fresh Nups by bringing them to interact with Nup153 and POM121. We speculate that this dual function of UBAP2L may bring together all required components at close vicinity of the NE, to ensure proper assembly of NPCs and their cellular function during early interphase (Fig. 10). Although our data are consistent with the role of UBAP2L in the biogenesis of new NPCs at the NE during early G1, at present, we cannot rigorously exclude the possibility that UBAP2L may also regulate a repair mechanism ensuring maintenance of the structural organization of the existing NPCs through its function on the Y-complex.

**Fig. 10.**
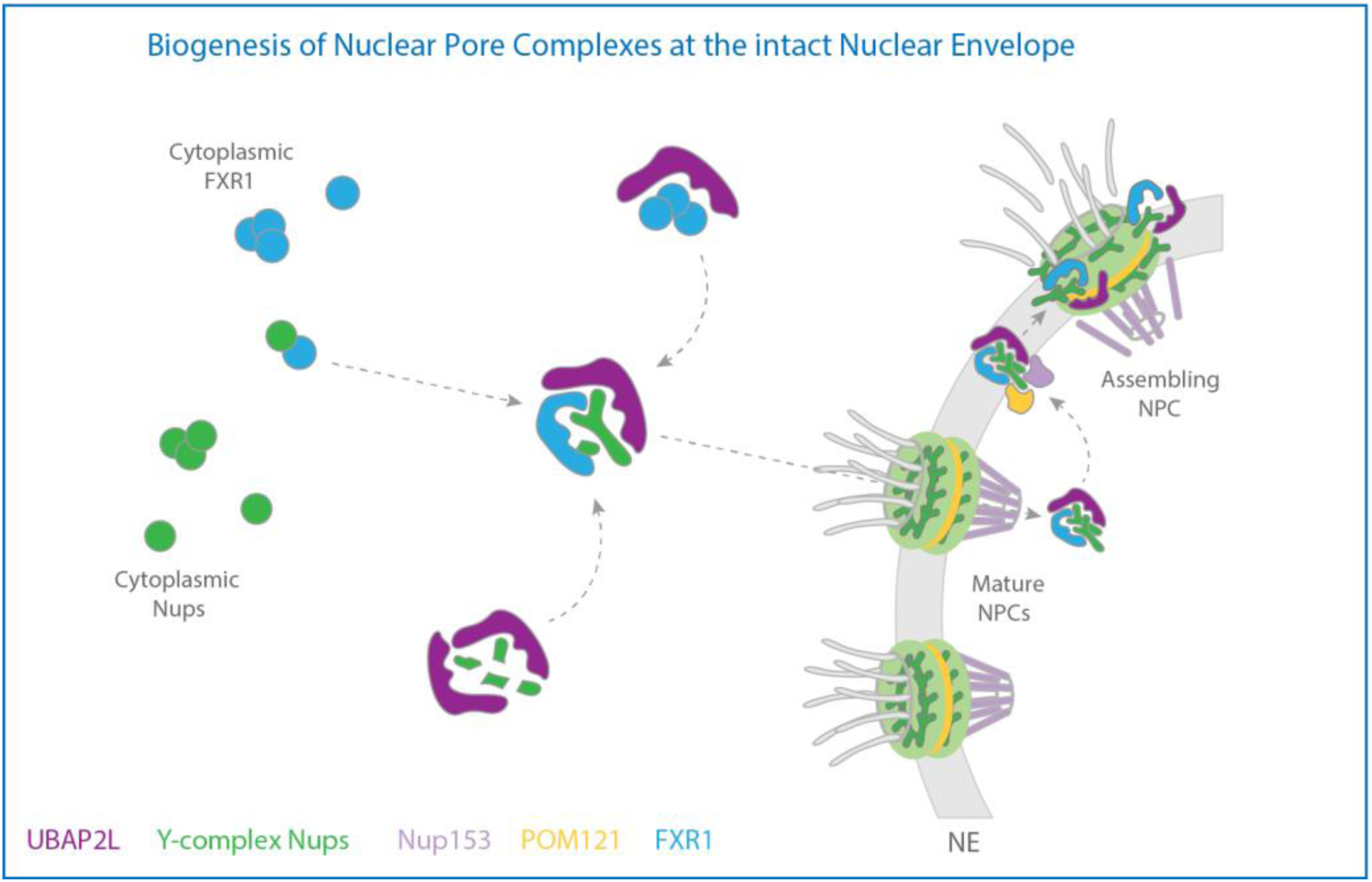
Hypothetical model how UBAP2L regulates the biogenesis of NPCs at the intact nuclear envelope. In the proximity of the nuclear envelope, UBAP2L (dark purple) interacts with cytoplasmic Y-complex nucleoporins (Nups) (green) and drives the formation of Y-complex. UBAP2L also interacts with the transporting factor of Nups in the cytoplasm, FXR1 (blue) and restricts its localization to NE during early G1 phase and ensures its interaction with Nups to fuel assembly and/or repair of NPCs. UBAP2L mediates the interaction of Y-complex Nups with Nup153 (light purple) and POM121 (yellow), which facilitates the assembly of functional and mature NPCs during interphase. This NPC biogenesis mechanism integrates the cytoplasmic and the nuclear NPC assembly signals and ensures efficient nuclear transport, adaptation to nutrient stress and cellular proliferative capacity, highlighting the importance of NPC homeostasis at the intact nuclear envelope.

The Y-complex named after its Y-shaped structure (Siniossoglou *et al*, 2000) is an essential component of the scaffold forming the cytoplasmic and the nuclear rings, respectively that encompass the inner ring of the NPC (von Appen *et al*, 2015). These sub-complexes oligomerize head to tail in a double-ring arrangement in each cytoplasmic and nuclear outer rings adding to 32 Y-complexes present in the human NPC (Bui *et al*, 2013). The molecular mechanisms governing the spatiotemporal assembly of the Y-complex and its oligomerization state into organized NPC scaffold remain uncharacterized. Our findings provide some insights into this biological riddle and identify UBAP2L as an important factor ensuring correct architectural organization of the NPC through the regulation of the formation of the Y-complex in human cells. Indeed, immunoprecipitation of Nup85 (Fig. 5G) and Nup96 (Fig. 5H) revealed interaction with other Y-complex Nups but this binding was reduced upon UBAP2L deletion. It remains to be determined if the oligomerization status of the Y-complex and its interaction with other NPC structural elements can be also regulated by UBAP2L.

In particular, the NPC cytoplasmic filaments component RanBP2 (also known as Nup358) was shown to wrap around the stems of Y-complexes to stabilize the scaffold (Huang *et al*, 2020b; von Appen *et al*, 2015). Similarly, the inner ring elements Nup188 and/or Nup205 were shown to contact Y-complex rings in vertebrate cells (Bui *et al*, 2013; Huang *et al*, 2020b; Kosinski *et al*, 2016; Lin *et al*, 2016). Future studies should provide further molecular insights into UBAP2L-mediated NPC assembly process and explain why the structure of the NPC visualized in our hands by super-resolution microscopy appears to be altered in the absence of UBAP2L (Fig. 5, A and D). Indeed, UBAP2L KO displayed moderately reduced number of NPC assemblies with mature symmetrical organization (Fig. 5, A and D) and the density of NPCs at the NE was decreased upon deletion of UBAP2L (Fig. 5, A and B, E and F). Because pre-pore structures observed by electron microscopy display eightfold arrangement already during early steps of NPC assembly by an inside-out extrusion mechanism at the intact NE (Otsuka *et al*, 2016), our results suggest that UBAP2L may act during initial steps of nuclear pore formation, prior to the described extrusion process. In agreement with this assumption, our data show that a portion of UBAP2L protein shuttles to the nucleus (Fig. S1, A to D) and that it can both localize to the NE (Fig. 1, A to C, and Fig. S2, A and B) and to the NPCs at the NE (Fig. 1, D to F) as well as it can interact with several Nups (Fig. 2, A to C, and Fig. 6A).

Moreover, the splitSMLM analysis suggested that UBAP2L appears to be more frequently localized at the nuclear ring labelled by Nup96 (Fig. 1, D and F) relative to the cytoplasmic ring, suggesting that UBAP2L may be transported through the existing mature NPCs to help the assembly of new pre-pore structures from the nuclear side. Future studies using electron microscopy (EM) approaches could shed some light on the presence of NPC assembly intermediates and on their structural organization in UBAP2L KO cells. Interestingly, interphase NPC assembly is initiated by the upstream Nups POM121 and Nup153 (Funakoshi *et al*, 2011; Vollmer *et al*, 2015) which recruit the Y-complex to the NE and their interaction with Y-complex is reduced in the absence of UBAP2L (Fig. 5, G and H). Thus, UBAP2L may not only regulate the formation of the Y-complex but also its timely recruitment to the NE through the binding to Nup153 and POM121.

Even though Nup153 and POM121 represent established upstream signals for the NPC biogenesis during interphase but not in the post-mitotic pathway, it was important to confirm that regulation of Nups by UBAP2L does not occur during mitotic exit, in particular in view of our recent analysis of the function of UBAP2L on PLK1 (Guerber *et al*, 2023). Indeed, we could also observe Nup defects in cells arrested in G1 phase (Fig. S2, C and D, G and H) and PLK1 inhibition could not reverse the Nup localization defects (Fig. S10, A and B), arguing that UBAP2L-dependent regulation of Nups could be largely uncoupled from its role in mitotic progression. Importantly, the described Nup localization defects upon deletion of UBAP2L could be first observed during late telophase/early G1 and throughout interphase but not in cells undergoing anaphase and early telophase (Fig. 4, A to F). In line with these observations, the regulation of FXR1 localization by UBAP2L appeared to be the most evident in early G1 compared to other cell cycle stages analyzed (Fig. S10, F to I). Thus, UBAP2L may regulate NPC assembly solely during interphase but not during mitotic exit.

A third pathway of the NPC assembly has been described in cells with rapid cell cycles and is based on the existence of the cytoplasmic stacks of double membranes, termed annulate lamellae (AL), which are structures containing partly assembled NPCs embedded in the endoplasmic reticulum (ER) membrane sheets, a feature associated with disturbances in NPC biogenesis (Hampoelz *et al*, 2016). AL can be inserted *en bloc* into the expanding NE in fly embryos (Hampoelz *et al*, 2016) and in higher eukaryotic cells, AL-based NPC assembly may represent an intermediate step in the postmitotic pathway (Ren *et al*, 2019). Interestingly, the splitSMLM analysis occasionally revealed the presence of linearly organized cytoplasmic assemblies of Nups in the absence of UBAP2L where RanBP2 was distributed symmetrically (Fig. S3A) and the cytoplasmic Nup foci induced by UBAP2L deletion did not contain Nup153 (Fig. 3A). These two observations are similar to the reported features of AL-NPC (Hampoelz *et al*, 2016) and could indicate that UBAP2L may, at least partially, contribute to the assembly of AL-NPC which will require further experimental efforts in the future.

Importantly, the biological significance of the UBAP2L-mediated assembly of the NPCs at the intact NE during early interphase is documented by defects in nucleocytoplasmic transport (Fig. 8) as well as by reduced proliferation capacity (Fig. S12, A to G) observed in UBAP2L-deficent cells. Although future studies will need to address whether UBAP2L-dependent regulation of NPC assembly can directly promote cell survival, UBAP2L was suggested to act as an oncogene (Chai *et al*, 2016; He *et al*, 2018; Li & Huang, 2014; Ye *et al*, 2017; Zhao *et al*, 2015; Guerber *et al*, 2022), and one could speculate that the role of UBAP2L in NPC biogenesis may explain, at least to some extent, the oncogenic potential of UBAP2L. This role of UBAP2L might be further regulated to meet differential demands on NPC functionality which may operate during changing cellular conditions such as stress or nutrient availability. Interestingly, deletion of Y-complex Nups can selectively affect survival and proliferation of colon cancer cells in response to presence of nutrients (Sakuma *et al*, 2020) and UBAP2L is sufficient to restore the NPC density after nutrient deprivation (Fig. 9), suggesting that UBAP2L-Nup pathway plays an important role under nutrient stress conditions which have been previously implicated in the regulation of NPC numbers in fission yeast (Varberg *et al*, 2021).

Taken together, our findings identify a molecular pathway driving biogenesis of mature and functional NPCs through spatiotemporal assembly of the Y-complex at the intact envelope in proliferating human cells. Our data further suggest a detailed molecular mechanism fueling the assembly of cytoplasmic Nups into Y-complex at the NE through a regulation of FXR1 protein by UBAP2L. UBAP2L strongly interacts with FXR1 (Fig. 2, B and C, and Fig. 6A, and Fig. S8, B to D) and regulates its localization to the NE (Fig. 7, and Fig. S5, A to G) and its interaction with the Y-complex (Fig. 5, G and H, and Fig. S4H) and the components of the dynein complex dynactin p150^Glued^ and BICD2 (Fig. 5, G and H, and Fig. S4H). Therefore, it can be speculated that UBAP2L may remodel cytoplasmic FXR1-assemblies found in mitotic cells specifically during early interphase and to promote the reported transport function of FXR1 by a minus-end directed microtubule-based mechanism towards NE (Agote-Aran *et al*, 2020). Thus, cells deficient for UBAP2L display both the cytoplasmic protein assemblies containing FXR1 or Nups, as a likely result of their assembly defects at the NE. How UBAP2L can execute its “chaperone-like” function on either FXR1 or Y-complex Nups and what are the regulatory mechanisms upstream of UBAP2L that may restrict its role to early interphase stage remain the subjects of future investigations. Interestingly, our data demonstrate that a region comprising 19 arginines present within the RGG domain may mediate UBAP2L’s function on FXR1 and Nups (Fig. 6). Globally, this mechanism appears to operate in an RNA-independent manner (Fig. S8, D and E), however, at this stage of analysis it cannot be excluded that specific RNAs might be involved in the UBAP2L-dependent regulation of Nups and FXR1. Consistent with a previous report (Huang *et al*, 2020a), mutation of the 19 arginines to alanines also led to loss of the asymmetric dimethylarginine (ADMA) signal (Fig. 6A). This raises an intriguing possibility, to be analyzed in the future, that ADMA or other arginine modifications of UBAP2L may regulate its function on Nups and their assembly into functional NPCs at the NE during early interphase.

## Materials and Methods

### Antibodies

The following primary antibodies were used in this study: rabbit anti-UBAP2L (1-430 aa) (Antibody facility IGBMC), mouse anti-FMRP (Antibody facility IGBMC), mouse anti-FXR1+2 (Antibody facility IGBMC), mouse monoclonal anti-GFP (Antibody facility IGBMC), mouse monoclonal anti-β-Actin (Sigma, A2228), rabbit polyclonal anti-GAPDH (Sigma, G9545), mouse monoclonal anti-α-Tubulin (Sigma, T9026), mouse monoclonal anti-FLAG® M2 (Sigma, F1804), rabbit polyclonal anti-FLAG® (Sigma, F7425), rabbit polyclonal anti-FXR1 (Sigma, HPA018246), rabbit polyclonal anti-Lamin A (C-terminal) (Sigma, L1293), rabbit monoclonal anti-Nup98 (C39A3) (Cell Signaling Technology, 2598), rabbit polyclonal anti-PRMT1 (A33) (Cell Signaling Technology, 2449), rabbit polyclonal anti-Tubulin (Abcam, ab18251), rabbit polyclonal anti-GFP (Abcam, ab290), rabbit polyclonal anti-FMRP (Abcam, ab17722), rabbit polyclonal anti-UBAP2L (Abcam, ab138309), rabbit monoclonal anti-Nup133 (Abcam, ab155990), mouse monoclonal anti-Nuclear Pore Complex Proteins (mAb414) (Abcam, ab24609), rabbit polyclonal anti-Nup153 (Abcam, ab84872), rabbit polyclonal anti-Nup188 (Abcam, ab86601), rabbit polyclonal anti-RanBP2 (Abcam, ab64276), rabbit polyclonal anti-Lamin B1 (Abcam, ab16048), rabbit monoclonal anti-NTF97/Importin beta (Abcam, ab2811), mouse monoclonal anti-Cyclin B1 (G-11) (Santa Cruz Biotechnology, sc-166757), mouse monoclonal anti-Cyclin E (HE12) (Santa Cruz Biotechnology, sc-247), mouse monoclonal anti-Nup133 (E-6) (Santa Cruz Biotechnology, sc-376763 AF488), mouse monoclonal anti-TIA-1 (Santa Cruz Biotechnology, sc-166247), mouse monoclonal anti-FXR1 (Millipore, 03-176), rabbit polyclonal anti-dimethyl-Arginine, asymmetric (ASYM24) (Millipore, 07-414), mouse monoclonal anti-Mps1 (Millipore, 05-682), rabbit polyclonal anti-G3BP1 (GeneTex, GTX112191), rabbit polyclonal anti-POM121 (GeneTex, GTX102128), rabbit polyclonal anti-Cyclin B1 (GeneTex, GTX100911), rabbit polyclonal anti-FXR2 (Proteintech, 12552-1-AP), mouse monoclonal anti-Nucleoporin p62 (BD Biosciences, 610497), mouse monoclonal anti-Ran (BD Biosciences, 610340), rabbit monoclonal anti-SEC13 (R&D systems, MAB9055), rabbit polyclonal anti-UBAP2L (1025-1087 aa) (Bethyl, A300-534A), rabbit polyclonal anti-Nup85 (Bethyl, A303-977A), rabbit polyclonal anti-Nup160 (Bethyl, A301-790A), rabbit polyclonal anti-CRM1/Exportin 1 (Novus, NB100-79802) and rat monoclonal anti-GFP (3H9) (ChromoTek, 3h9-100), rabbit polyclonal anti-LC3B (Novus biological, NB100-2331), guinea pig polyclonal anti-p62 (Progen, GP62-C), rabbit monoclonal anti-DNAJB6 (Abcam, ab198995), rabbit monoclonal anti-BiCD2 (Sigma, HPA023013), mouse monoclonal anti-p150^Glued^ (BD biosciences, 610473).

Secondary antibodies used were the following: goat polyclonal anti-Mouse CF680 (Sigma, SAB4600199), goat polyclonal anti-Chicken CF660C (Sigma, SAB4600458), goat polyclonal anti-Mouse AF647 (Thermo Fisher Scientific, A-21236), goat polyclonal anti-Mouse AF568 (Thermo Fisher Scientific, A-11031), goat polyclonal anti-Mouse AF555 (Thermo Fisher Scientific, A-11029), goat polyclonal anti-Mouse AF488 (Thermo Fisher Scientific, A-21424), goat polyclonal anti-Rabbit AF647 (Thermo Fisher Scientific, A-21245), goat polyclonal anti-Rabbit AF568 (Thermo Fisher Scientific, A-11036), goat polyclonal anti-Rabbit AF555 (Thermo Fisher Scientific, A-21429), goat polyclonal anti-Rabbit AF488 (Thermo Fisher Scientific, A-11034), goat Anti-Mouse IgG antibody (HRP) (GeneTex, GTX213111-01), goat Anti-Mouse IgG antibody (HRP) (GeneTex, GTX213110-01) and goat Anti-Rat IgG antibody (HRP) (Cell Signaling Technology, 7077S).

### Generation of UBAP2L KO cell lines

UBAP2L knock-out (KO) in HeLa cells were described previously (Guerber *et al*, 2023). UBAP2L KO in Nup96-GFP knock-in (KI) U2OS (CLS Cell Line Service, 300174; a generous gift of Arnaud Poterszman, IGBMC) cell lines were generated using CRISPR/Cas9 genome editing system as described previously (Jerabkova *et al*, 2020). Two guide RNAs (gRNA) were designed using the online software Benchling (https://www.benchling.com/), 5’-TGGCCAGACGGAATCCAATG-3’ and 5’-GTGGTGGGCCACCAAGACGG-3’, and cloned into pX330-P2A-EGFP/RFP (Zhang *et al*, 2017) through ligation using T4 ligase (New England Biolabs). Nup96-GFP KI U2OS cells were transfected using X-tremeGENE^TM^ 9 DNA Transfection Reagent (Roche), and 24h after transfection, GFP and RFP double positive cells were collected by FACS (BD FACS Aria II), cultured for 2 days and seeded with FACS into 96-well plates. Obtained UBAP2L KO single-cell clones were validated by Western blot and sequencing of PCR-amplified targeted fragment by Sanger sequencing (GATC). The following primers were used for PCR amplification: 5′-TGCTGAGTGGAGAATGGTTA-3′ (forward) and 5′-AGACTGGTGGCAGTTGGTAG-3′ (reverse). Primers used for cloning and sequencing are described in Table S1.

### Cell culture

All cell lines were cultured at 37°C in 5% CO2 humidified incubator. HeLa (Kyoto) and its derived UBAP2L KO cell lines were cultured in Dulbecco’s Modified Eagle Medium (DMEM) (4.5 g/L glucose) supplemented with 10% fetal calf serum (FCS), 1% Penicillin + 1% Streptomycin. U2OS were cultured in DMEM (1 g/L glucose) supplemented with 10% FCS + Gentamicin 40 µg/mL. Nup96-GFP KI U2OS and its derived UBAP2L KO cell lines were cultured in DMEM (1 g/L glucose) supplemented with 10% FCS, Non-Essential Amino Acids + Sodium Pyruvate 1 mM + Gentamicin 40 µg/mL.

### Cell cycle synchronization treatments

Cells were synchronized in different stages of cell cycle by double thymidine block and release (DTBR) protocol. Briefly, cells were treated with 2 mM thymidine for 16h, washed out (three times with warm thymidine-free medium), then released in fresh thymidine-free culture medium for 8h, treated with 2 mM thymidine for 16h again, washed out, and then released in fresh thymidine-free culture medium for different time periods (0, 3, 6, 8, 9, 10 and 12h). 0h time point corresponds to G1/S phase, approximately 8h to 9h to mitotic peak, 10h to mitotic exit and 12h to early G1 phase. Cells were synchronized in G1 phase using lovastatin for 16h at 10 µM final concentration and in G0/G1 using Psoralidin (3,9-Dihydroxy-2-prenylcoumestan) for 24h at 5 µM final concentration. Cells were synchronized in prometaphase using Nocodazole for 16h at 100 ng/ml, STLC for 16h at 5 µM, and monastrol for 16h at 100 µM final concentration.

### Plasmids

All pEGFP-C1-UBAP2L wild type (WT) (NCBI, NM_014847.4), pEGFP-C1-UBAP2L UBA (1-97 aa), pEGFP-C1-UBAP2L ΔUBA (Δ1-97 aa), pEGFP-C1-UBAP2L 98-430 aa, pEGFP-C1-UBAP2L 1-430 aa, pEGFP-C1-UBAP2L Δ1-429 aa, pEGFP-C1-UBAP2L Δ(UBA+RGG) (Δ1-195 aa) and pEGFP-C1-FXR1 WT (NCBI, NM_001013438.3) plasmids were generated by Stephane Schmucker (IGBMC). pcDNA3.1-Flag-N-UBAP2L WT (NCBI, NM_014847.4) was generated by Evanthia Pangou (IGBMC). Primers used for cloning are described in Table S1. pEGFP-C1 was purchased from Clontech. pcDNA3.1-Flag-N was obtained from IGBMC cloning facility, and pcDNA3.1-Flag-UBAP2L R131-190A was a generous gift of Zhenguo Chen (Southern Medical University, P. R. China) (Huang *et al*, 2020a). pEGFP-C1-Nup85 was kindly provided by Valérie Doye (Institut Jacques Monod, Paris), and pXRGG-GFP was kindly provided by Jan M. van Deursen (Hamada *et al*, 2011; Love *et al*, 1998).

### Plasmid and siRNA transfections

Lipofectamine 2000 (Invitrogen), jetPEI-DNA transfection reagent (Polyplus-transfection) and X-tremeGENE^TM^ 9 DNA Transfection Reagent (Roche) were used to perform plasmid transient transfection according to the manufacturer’s instructions. Lipofectamine RNAiMAX (Invitrogen) was used to deliver siRNAs for gene knock-down (KD) according to the manufacturer’s instructions at a final concentration of 20 to 40 nM siRNA. The following siRNA oligonucleotides were used: Non-targeting individual siRNA-2 5’-UAAGGCUAUGAAGAGAUAC-3’ (Dharmacon), UBAP2L siRNA 5’-CAACACAGCAGCACGUUAU-3’ (Dharmacon) and FXR1 siRNA-1 5’-AAACGGAAUCUGAGCGUAA-3’ (Dharmacon).

### Protein preparation and Western blotting

Cells were collected by centrifugation at 200 g for 4 min at 4 °C and washed twice with cold phosphate buffered saline (PBS), and cell lysates for Western blot were prepared using 1X RIPA buffer (50 mM Tris-HCl pH 7.5, 150 mM NaCl, 1% Triton X-100, 1 mM EDTA, 1 mM EGTA, 2 mM Sodium pyrophosphate, 1 mM Na3VO4 and 1 mM NaF) supplemented with protease inhibitor cocktail (Roche) and incubated on ice for 30 min. After centrifugation at 16 000 g for 15 min at 4 °C, cleared supernatant was transferred to the new clean Eppendorf tubes and total protein concentration was measured using Bradford assay by Bio-Rad Protein Assay kit (Bio-Rad). Nuclear and cytoplasmic proteins were prepared using the NE-PER nuclear and cytoplasmic extraction reagent kit (Thermo Scientific™, 78833). Protein samples were boiled for 8 min at 95 °C in 1X Laemmli buffer (LB) with β-Mercaptoethanol (BioRad, 1610747), resolved on 10% polyacrylamide gels or pre-cast 4-12% Bis-Tris gradient gels (Thermo Scientific, NW04120BOX) or pre-cast NuPAGE™ 3-8% Tris-Acetate gradient Gels (Thermo Scientific, EA0378BOX) and transferred to a polyvinylidene difluoride (PVDF) membrane (Millipore, IPFL00010) using semi-dry transfer unit (Amersham) or wet transfer modules (BIO-RAD Mini-PROTEAN® Tetra System). Membranes were blocked in 5% non-fat milk powder, 5% bovine serum albumin (BSA, Millipore, 160069), or 5% non-fat milk powder mixed with 3% BSA and resuspended in TBS-T (Tris-buffered saline-T: 25 mM Tris-HCl, pH 7.5, 150 mM NaCl 0.05% Tween) for 1h at room temperature, followed by incubation with antibodies diluted in TBS-T 5% BSA/5% milk. All incubations with primary antibodies were performed for overnight at 4°C. TBS-T was used for washing the membranes. Membranes were developed using SuperSignal West Pico (Pierce, Ref. 34580) or Luminata Forte Western HRP substrate (Merck Millipore, Ref. WBLUF0500).

### Immunoprecipitations

Cell lysates for immunoprecipitations (IP) were prepared using 1X RIPA buffer supplemented with protease inhibitor cocktail and incubated on ice for 1h. When indicated, 1X RIPA buffer was supplemented with with RNase A or Benzonase. After centrifugation at 16 000 g for 15 min at 4 °C, cleared supernatant was transferred to the new clean Eppendorf tubes. Lysates were equilibrated to volume and concentration.

For endogenous IP experiments, IgG and target specific antibodies as well as protein G sepharose 4 Fast Flow beads (GE Healthcare Life Sciences) were used. Samples were incubated with the IgG and target specific antibodies overnight at 4 °C with rotation. Beads were blocked with 3% BSA diluted in 1X RIPA buffer and incubated for 2h at 4 °C with rotation. Next, the IgG/ specific antibodies-samples and blocked beads were incubated in 1.5 ml Eppendorf tubes to a final volume of 1 ml 4h at 4 °C with rotation. The incubated IgG/ specific antibodies-samples-beads were washed with washing buffer (25 mM Tris-HCl pH 7.5, 300 mM NaCl, 0.5% Triton X-100, 0.5 mM EDTA, 0.5 mM EGTA, 1 mM Sodium pyrophosphate, 0.5 mM Na3VO4 and 0.5 mM NaF) or TBS-T supplemented with protease inhibitor cocktail 4 to 6 times for 10 min each at 4°C with rotation. Beads were pelleted by centrifugation at 200 g for 3 min at 4 °C. The washed beads were directly eluted in 2X LB with β-Mercaptoethanol and boiled for 12 min at 95 °C for Western blot.

For GFP-IP/Flag-IP experiments, GFP-Trap A agarose beads (Chromotek) or Flag beads (Sigma) were used. Cells expressing GFP-or Flag-tagged plasmids for at least 24 h were used to isolate proteins using 1X RIPA buffer supplemented with protease inhibitor cocktail. Beads were blocked with 3% BSA diluted in 1X RIPA buffer and incubated for 2h at 4 ⁰C with rotation. Samples were incubated with the blocked beads for 2h or overnight at 4 °C with rotation, and the beads were washed and boiled as for endogenous IP.

### Immunofluorescence

Cells grown on glass coverslips (Menzel-Glaser) were washed twice in PBS and then fixed with 4% paraformaldehyde (PFA, Electron Microscopy Sciences 15710) in PBS for 15 min at room temperature, washed 3 times for 5 min in PBS and permeabilized with 0.5% NP-40 (Sigma) in PBS for 5 min. Cells were washed 3 times for 5 min in PBS and blocked with 3% BSA in PBS-Triton 0.01% (Triton X-100, Sigma, T8787) for 1h. Cells were subsequently incubated with primary antibodies in blocking buffer (3% BSA in PBS-Triton 0.01%) for 1h at room temperature, washed 3 times for 8 min in PBS-Triton 0.01% with rocking and incubated with secondary antibodies in blocking buffer for 1h at room temperature in the dark. After incubation, cells were washed 3 times for 8 min in PBS-Triton 0.01% with rocking in the dark and glass coverslips were mounted on glass slides using Mowiol containing 0.75 μg/ml DAPI (Calbiochem) and imaged with a 100x or 63x objective using Zeiss epifluorescence microscope. For mitotic cells immunofluorescence, cells were collected from dishes with cell scrapers, centrifuged on Thermo Scientific Shandon Cytospin 4 Cytocentrifuge for 5 min at 1000 rpm and fixed immediately with 4% PFA for 15 min at room temperature.

For nucleoporins (Nups) immunofluorescence, cells grown on glass coverslips were washed twice in PBS and then fixed with 1% PFA in PBS for 10 min at room temperature, washed 3 times for 5 min in PBS and permeabilized with 0.1% Triton X-100 and 0.02% SDS (Euromedex, EU0660) in PBS for 5 min. After permeabilization, cells were washed 3 times for 5 min in PBS and blocked with 3% BSA in PBS-Triton 0.01% for 1h at room temperature or overnight at 4 °C. Cells were subsequently incubated with primary antibodies in blocking buffer (3% BSA in PBS-T) for 1h at room temperature, washed 3 times for 8 min with rocking in blocking buffer and then incubated with secondary antibodies in blocking buffer for 1h at room temperature in the dark. After incubation, cells were washed 3 times for 8 min with rocking in blocking buffer in the dark and then permeabilized again with 0.1% Triton X-100 and 0.02% SDS in PBS for 1 min and post-fixed for 10 min with 1% PFA in PBS at room temperature in the dark. Then coverslips were washed twice in PBS for 5 min and mounted on glass slides using Mowiol containing 0.75 μg/ml DAPI.

An adapted protocol was used for the experiments presented in Fig. S2A as described previously (Guerber *et al*, 2023). After the appropriate synchronization using DTBR, the cytoplasm was extracted from the cells to remove the large cytoplasmic fraction of UBAP2L by incubating the coverslips in cold 0,01% Triton X-100 for 90sec. 4% PFA was immediately added to the coverslips after the pre-extraction and the standard IF protocol was followed.

### Sample preparation for single molecule localization microscopy

For super-resolution single molecule localization microscopy (splitSMLM), cells were plated on 35 mm glass bottom dish with 14 mm micro-well #1.5 cover glass (Cellvis). Cells were washed twice with PBS (2 ml/well) and then fixed with 1% PFA in PBS for 15 min at room temperature, washed 3 times for 5 min in PBS (store samples submerged in PBS at 4 °C until use) and permeabilized with 0.1% Triton X-100 (Tx) in PBS (PBS/Tx) for 15 min. Cells were blocked with 3% BSA in 0.1% PBS/Tx (PBS/Tx/B) for 1h and then incubated with primary antibodies (optimal working concentration of primary antibody is 2 μg/ml) in PBS/Tx/B (200 μl/well) for overnight at 4 °C in wet chamber. After incubation, cells were washed 3 times for 8 min with rocking in PBS/Tx/B and subsequently incubated with secondary antibodies (optimal working concentration of secondary antibody is 4 μg/ml) in PBS/Tx/B (200 μl/well) for 2h at room temperature in the dark. Immediately after, cells were washed 3 times for 8 min with rocking in PBS/Tx and post-fixed for 10 min with 1% PFA at room temperature in the dark, then cells were washed twice in PBS and kept in PBS in the dark.

The samples were imaged in a water-based buffer that contained 200 U/ml glucose oxidase, 1000 U/ml catalase, 10% w/v glucose, 200 mM Tris-HCl pH 8.0, 10 mM NaCl and 50 mM MEA. 2 mM cyclooctatetraene was added to the buffer for multi-color imaging (Andronov *et al*, 2022). The mixture of 4 kU/ml glucose oxidase (G2133, Sigma) and 20 kU/ml catalase (C1345, Sigma) was stored at −20 °C in an aqueous buffer containing 25 mM KCl, 4 mM TCEP, 50% v/v glycerol and 22 mM Tris-HCl pH 7.0. MEA-HCl (30080, Sigma) was stored at a concentration of 1M in H2O at −20 °C. Cyclooctatetraene (138924, Sigma) was stored at 200 mM in dimethyl sulfoxide at −20 °C. The samples were mounted immediately prior to imaging filling the cavity of the glass-bottom petri dishes with ∼200 µl of the imaging buffer and placing a clean coverslip on top of it, which allowed imaging for ≥ 8 hours without degradation of the buffer. After imaging, the samples were washed once with PBS and kept in PBS at +4 °C.

### Single molecule localization microscopy

The SMLM experiments were performed on a splitSMLM system (Andronov *et al*, 2022) that consisted of a Leica DMI6000B microscope; an HCX PL APO 160x/1.43 Oil CORR TIRF PIFOC objective; a 642 nm 500 mW fiber laser (MBP Communication Inc.) for fluorescence excitation and a 405 nm 50 mW diode laser (Coherent Inc.) for reactivation of fluorophores.

The sample was illuminated through a Semrock FF545/650-Di01 dichroic mirror and the fluorescence was filtered with Semrock BLP01-532R and Chroma ZET635NF emission filters. For single-color imaging that was used for estimation of the NPC density at the NE, the fluorescence was additionally filtered with a Semrock BLP01-635R-25 long-pass filter and was projected onto an Andor iXon+ (DU-897D-C00-#BV) EMCCD camera.

For multi-color imaging, the fluorescence was split into two channels with a Chroma T690LPXXR dichroic mirror inside an Optosplit II (Cairn Research) image splitter. The short-wavelength channel was additionally filtered with a Chroma ET685/70m bandpass filter and both channels were projected side-by-side onto an Andor iXon Ultra 897 (DU-897U-CS0-#BV) EMCCD camera.

The SMLM acquisitions began with a pumping phase, during which the sample was illuminated with the 642 nm laser but the fluorescence was not recorded due to a very high density of fluorophores in a bright state. When the density dropped to a level that allowed observation of individual molecules, the images started to be recorded. Pumping and imaging were performed at 30-50% of maximal power of the 642 nm laser. When the density of fluorophores in the bright state dropped further due to photobleaching, the sample started to be illuminated with the 405 nm laser for reactivation of fluorophores. The intensity of the 405 nm laser was increased gradually to account for the photobleaching. For estimation of the NPC density, to increase speed, the pumping and imaging were performed at 100% laser power and the acquisitions were stopped after about 2 min of imaging.

### Processing of single molecule localization microscopy data

The fitting of single-molecule localizations was done in the Leica LAS X software with the “direct fit” method. For single-color imaging, the obtained localization tables were corrected for drift and reconstructed as 2D histograms with a pixel size of 15 nm in SharpViSu (Andronov *et al*, 2016). For multi-color imaging, the localizations were first unmixed in SplitViSu (Andronov *et al*, 2022). Next, they were corrected for drift and for relocalizations in SharpViSu, and reconstructed as 2D histograms with a pixel size of 5 nm.

For quantification of the rotational symmetry of the NPCs, individual NPCs were picked manually on the NE of each imaged cell. Only particles that are in focus and in correct “top view” orientation were selected. For the analysis, the localizations within a radius of 130 nm from the manually picked center of each NPC were used. The obtained particles were aligned in smlm_datafusion2d with random rotation of every particle by n ⋅ 45°, n = [0, 7], after each alignment iteration (Heydarian *et al*, 2018). The aligned particles were then converted to polar coordinates and localizations with radii from 50 to 70 nm were kept for further analysis. A sine function with a period of π/4 was fitted to the polar angle distribution of the sum of all aligned particles. The localizations were split into eight sectors using the minima of the sine function as the edges of the sectors. The number of localizations within each sector was calculated for each NPC. For a given NPC, a sector was considered occupied if the number of localizations within it was higher than the half of the mean number of localizations per sector for this NPC. The quantified number of subunits of an NPC is the number of the occupied sectors.

The axial and radial profiles of the NPCs were obtained as described previously (Andronov *et al*, 2022). For radial profiles, the localizations of co-imaged proteins were transformed using the alignment parameters of Nup96 after 8-fold alignment in smlm_datafusion2d (Heydarian *et al*, 2018). For the “side view” profile, the axial profiles of individual particles were calculated in Fiji (Schindelin *et al*, 2012), averaging through the whole thickness of the NPC. The axial profiles of Nup96 particles were fitted with a sum of two Gaussians in Matlab. Nup96 particles and co-imaged proteins were aligned using this fit of Nup96.

### Live-cell imaging

For FXR1 mitotic granules assay, WT and UBAP2L KO HeLa cells expressing GFP-FXR1 were grown on 35/10 mm 4 compartment glass bottom dishes (Greiner Bio-One, 627871) and synchronized by double thymidine block, released for 8h and analyzed by Nikon PFS spinning disk (63× objective) for 9h. Z-stacks (7 μm range, 1 μm step) were acquired every 5 min and movies were made with maximum intensity projection images for every time point shown at speed of 7 frames per second. Image quantification analysis was performed using ImageJ software.

For protein import and export assay, WT and UBAP2L KO HeLa cells were grown on 8-well Chambered Coverglass w/non-removable wells (Thermo Fisher Scientific, 155411PK) and transfected with the reporter plasmid XRGG-GFP for 30h, and incubated with full media with SiR-DNA 1:1500 and Verapamil 1:1000 for at least 1h before filming. Then SiR-DNA and Verapamil were kept with media and cells were incubated in media with 0.01 μM dexamethasone. Dexamethasone-induced nuclear import of XRGG-GFP was recorded by Leica CSU-W1 spinning disk (63X objective) for 129 min (1 acquisition every 1 min, 12 μm range, 3 μm step). For nuclear export, dexamethasone was washed out at 129 min time point with warm dexamethasone-free medium, cells were incubated with full media with SiR-DNA 1:6000 and Verapamil 1:4000 and nuclear export of XRGG-GFP was recorded for 170 min (1 acquisition every 1 min, 12 μm range, 3 μm step). Image quantification analysis was performed using ImageJ software.

### Nuclear envelope intensity analysis of nucleoporins

A CellProfiler software pipeline was previously generated by Arantxa Agote-Aran (Agote-Aran *et al*, 2020) that automatically recognizes cell nuclei based on the DAPI fluorescent image. A threshold of nuclei size was applied to the pictures to exclude too small or too big nuclei and nuclei edges were enhanced using the Prewitt edge-finding method. This allowed identification and measurement of the nuclei area, form factor and nuclear mean intensity of desired channels. The parameters’ measurements of the software were exported to an Excel file and statistically analysed. At least 200 cells from three different biological replicates were measured.

### Colony formation assay

500 WT and UBAP2L KO HeLa cells were seeded per well in 6-well plates and incubated at 37 °C in 5% CO2 for 7 days until colonies formed. Cells were washed with 1X PBS, fixed with 4% PFA and stained with 0,1% Crystal Violet for 30 min. The number of colonies was first manually counted and then automatically quantified with Fiji software.

### Flow cytometry

For cell death analysis, HeLa cells were spun down and resuspended in cold PBS supplemented with 50 μg/ml propidium iodide (PI) (Sigma-Aldrich, Ref. P4170). PI positive cells were analyzed by BD FACS CelestaTM Flow Cytometer.

### Experimental design, data acquisition and statistical analysis

All experiments were done in a strictly double-blind manner. At least three independent biological replicates were performed for each experiment (unless otherwise indicated) and image quantifications were carried out in a blinded manner. Curves and graphs were made using GraphPad Prism and Adobe Illustrator software. Data was analyzed using one-sample two-tailed T-test or two sample two-tailed T-test (two-group comparison or folds increase relative to the control, respectively). A p-value less than 0.05 (typically ≤ 0.05) was considered statistically significant and stars were assigned as follows: *P < 0.05, **P < 0.01, ***P < 0.001, ****P < 0.0001. In all graphs, results were shown as mean ± SD, and details for each graph were listed in the corresponding figures’ legends.

## Acknowledgements

We thank the members of the I. Sumara and R. Ricci laboratories for helpful discussions on the manuscript. Y.L. was supported by a PhD fellowship from the China Scholarship Council (CSC) and postdoctoral fellowship from SATT Conectus Alsace. X.L., J.L., M.Q., and L.R. were supported by a PhD fellowship from the China Scholarship Council (CSC) and A.A.A., and L.G. were supported by Labex international PhD fellowship from IGBMC and IMC-Bio graduate school. E.P. was supported by postdoctoral fellowships from the “Foundation pour la recherché Médicale” (FRM) and ANR-10-LABX-0030-INRT. L.A., and B.P.K. acknowledge support by Institut National du Cancer (INCa) and by the French Infrastructure for Integrated Structural Biology (FRISBI) ANR-10-INSB-05-01, Instruct-ERIC and iNEXT-Discovery. Research in I.S. laboratory was supported by the grant ANR-10-LABX-0030-INRT, a French State fund managed by the Agence Nationale de la Recherche under the frame program Investissements d’Avenir ANR-10-IDEX-0002-02, IGBMC, CNRS, Fondation ARC pour la recherche sur le cancer, Institut National du Cancer (INCa), Agence Nationale de la Recherche (ANR), Ligue Nationale contre le Cancer, Sanofi iAward Europe and Programme Fédérateur Aviesan, Plan Cancer, National collaborative project: “NANOTUMOR”.

## Author contributions

Y.L., and L.A. designed and performed experiments and helped writing the manuscript. X.L., J.L., L.G., L.L., A.A.A., E.P., L.R., C.K., M.Q., S.S., and L.C. performed experiments. Z.Z. helped performing experiments. D.R., M.G., and B.P.K. helped designing the experiments and supervising. I.S. supervised the project, designed experiments and wrote the manuscript with input from all authors.

## Data and materials availability

All data needed to evaluate the conclusions in the paper are present in the paper and/or the Supplementary Materials.

## Supplementary Materials for

### Supplemental figures

**Fig. S1.**
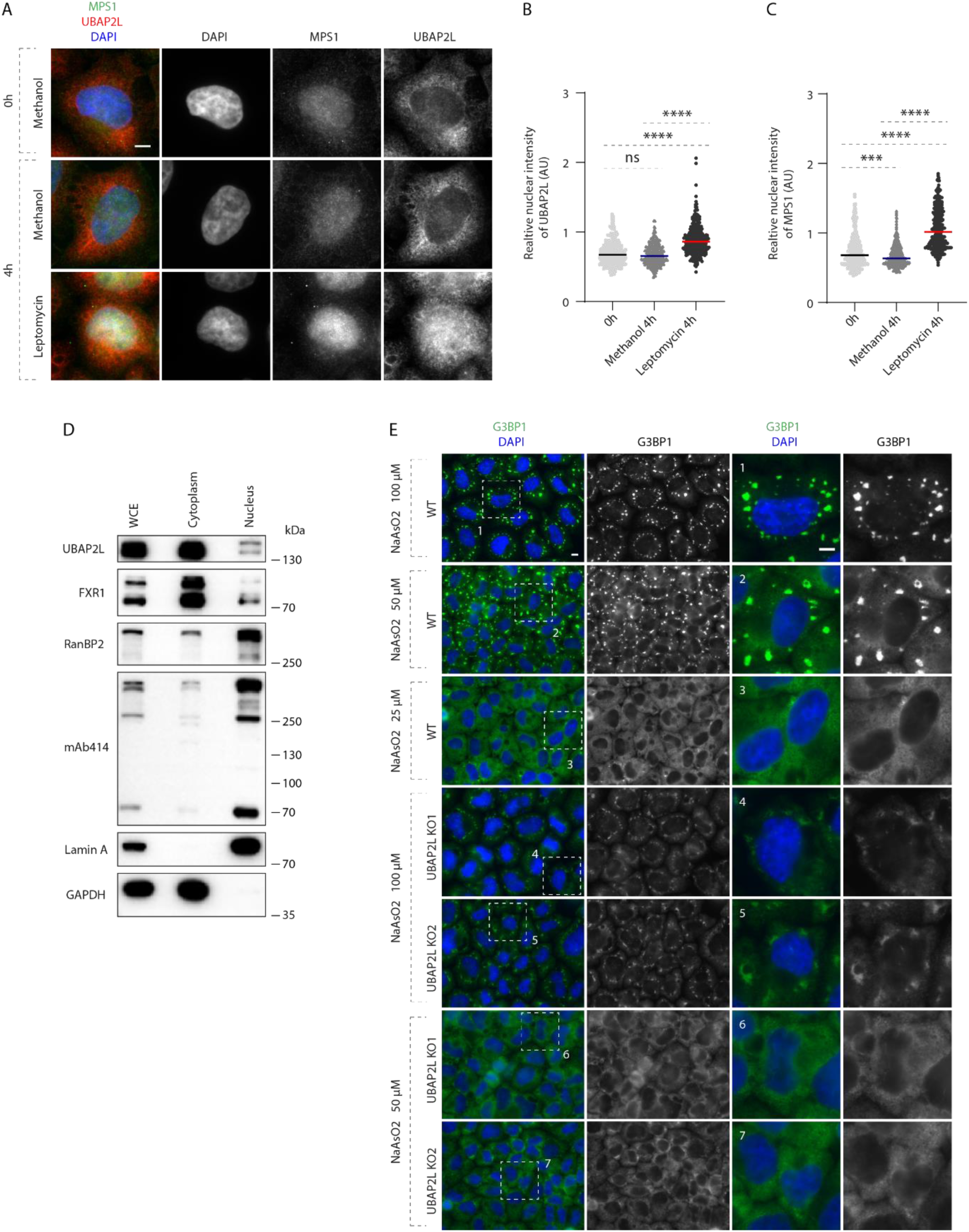
UBAP2L shuttles between cytoplasm and nucleus. (**A to C**) Representative immunofluorescence images depicting the cytoplasmic and nuclear localization of UBAP2L and MPS1 (also known as protein kinase TTK) after treatment with the Leptomycin B (inhibitor of nuclear export factor Exportin 1) (100 ng/ml) for 4h (**A**). Nuclei were stained with DAPI. The realtive nuclear intensity (AU) of UBAP2L (**B**) and MPS1 (**C**) shown in (**A**) was quantified. At least 150 cells per condition were analyzed (mean ± SD, ns: not significant, ***P < 0.001, ****P < 0.0001; two-tailed t-test, *N* = 3). Scale bar, 5 μm. **(D)** Protein levels of UBAP2L, FXR1 and Nups were analyzed by Western blot in the whole cell extract (WCE) and in nuclear and cytoplasmic fractions of HeLa cells. **(E)** Representative immunofluorescence images of WT and UBAP2L KO HeLa cells depicting formation of stress granules (SGs) labelled by G3BP1 at indicated arsenite concentrations. The magnified framed regions are shown in the corresponding numbered panels. Nuclei were stained with DAPI. Scale bars, 5 μm.

**Fig. S2.**
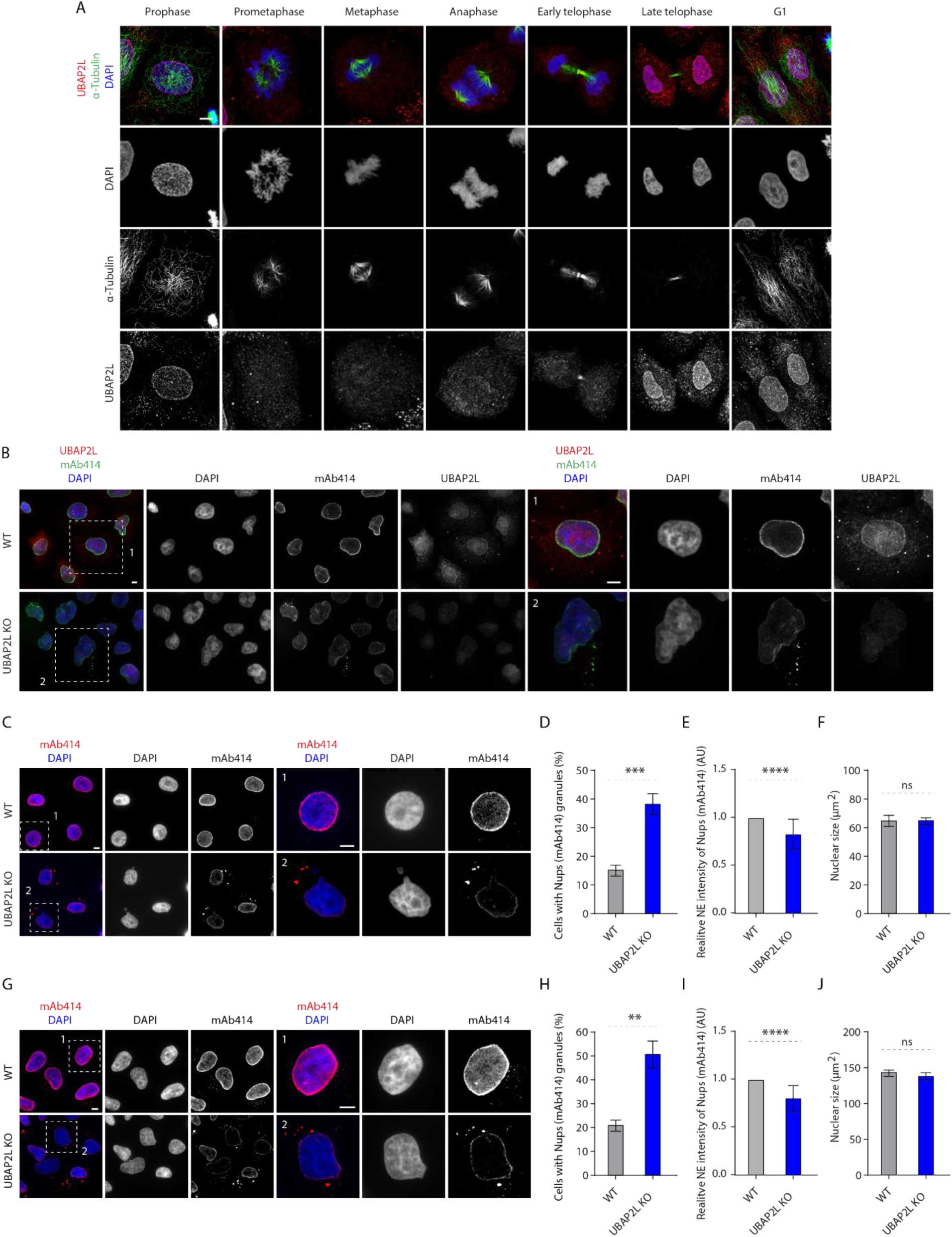
Localization of UBAP2L during cell cycle progression. **(A)** Representative immunofluorescence images depicting the localization of UBAP2L in HeLa cells after chemical pre-extraction of the cytoplasm using 0,01% of Triton X-100 for 90sec in indicated cell cycle stages and visualized by UBAP2L antibody. Nuclei and chromosomes were stained with DAPI. Scale bar, 5 μm. **(B)** Representative immunofluorescence images depicting the localization of nucleoporins (Nups) and UBAP2L in asynchronously proliferating wild type (WT) and UBAP2L Knock-out (KO) HeLa cells visualized by mAb414 and UBAP2L antibodies. Nuclei were stained with DAPI. The magnified framed regions are shown in the corresponding numbered panels. Note that UBAP2L signal is absent in UBAP2L-deleted cells. Scale bars, 5 μm (**C to F**) Representative immunofluorescence images depicting the localization and NE intensity of Nups (mAb414) and nuclear size in WT and UBAP2L KO HeLa cells synchronized in G1 phase by lovastatin (10 µM) for 16h (**C**). The magnified framed regions are shown in the corresponding numbered panels. Scale bars, 5 μm. The cells with Nups (mAb414) granules (**D**), the NE intensity of Nups (mAb414) (**E**) and the nuclear size (**F**) shown in (**C**) were quantified. At least 150 cells per condition were analyzed (mean ± SD, ns: not significant, ***P < 0.001, ****P < 0.0001, two-tailed t-test, *N* = 3) (**G to J**) Representative immunofluorescence images depicting the localization and NE intensity of Nups (mAb414) and nuclear size in WT and UBAP2L KO HeLa cells synchronized in G0/G1 phase by Psoralidin (5 µM) for 24h (**G**). The magnified framed regions are shown in the corresponding numbered panels. Scale bars, 5 μm. The cells with Nups (mAb414) granules (**H**), the NE intensity of Nups (mAb414) (**I**) and the nuclear size (**J**) shown in (**G**) were quantified. At least 200 cells per condition were analyzed (mean ± SD, ns: not significant, **P < 0.01, ****P < 0.0001, two-tailed t-test, *N* = 3).

**Fig. S3.**
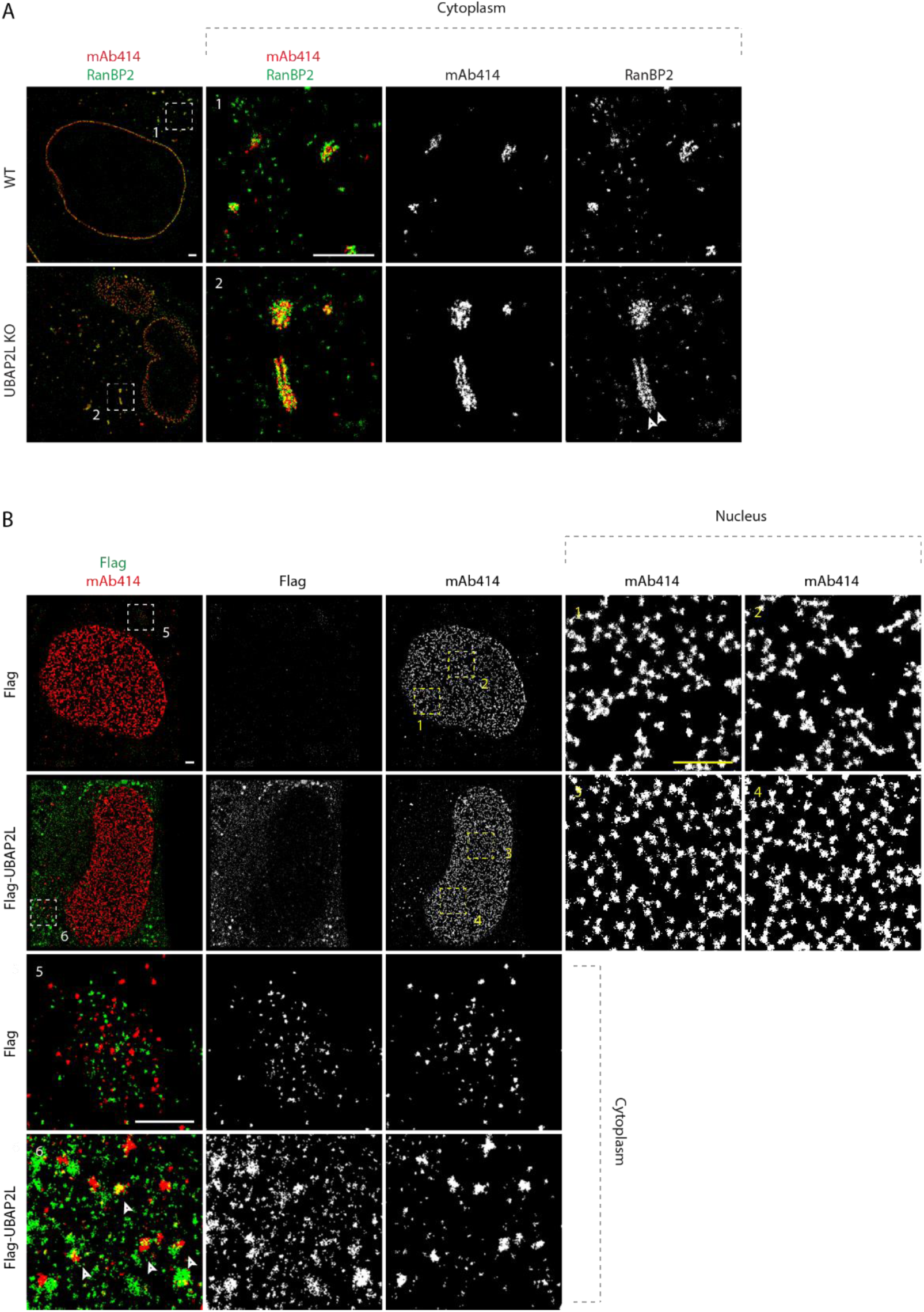
UBAP2L may inhibit formation of cytoplasmic annulate lamellae (AL) or AL-like Nup assemblies. **(A)** Representative splitSMLM immunofluorescence images depicting the localization of NPC components corresponding to central channel (FG-Nups labeled by mAb414) and cytoplasmic filaments (RanBP2) at NE and in the cytoplasm in WT and UBAP2L KO HeLa cells synchronized in interphase by DTBR at 12h. Note that unlike at the NE where RanBP2 can localize exclusively to the cytoplasmic side of the NPCs (Fig. 5A), deletion of UBAP2L leads to the accumulation of the Nup assemblies in the cytoplasm with a symmetric distribution of RanBP2. The magnified framed regions are shown in the corresponding numbered panels. Scale bars, 1000 and 300 nm, respectively. **(B)** Representative SMLM immunofluorescence images of FG-Nups (mAb414) at the nuclear surface in interphase HeLa cells expressing Flag alone or Flag-UBAP2L for 35h and synchronized by DTBR at 12h. The magnified framed regions are shown in the corresponding numbered panels and corresponding quantification is shown in Fig. 5C. The arrowheads indicate the cytoplasmic colocalization of UBAP2L and FG-Nups. Scale bars, 1000 nm.

**Fig. S4.**
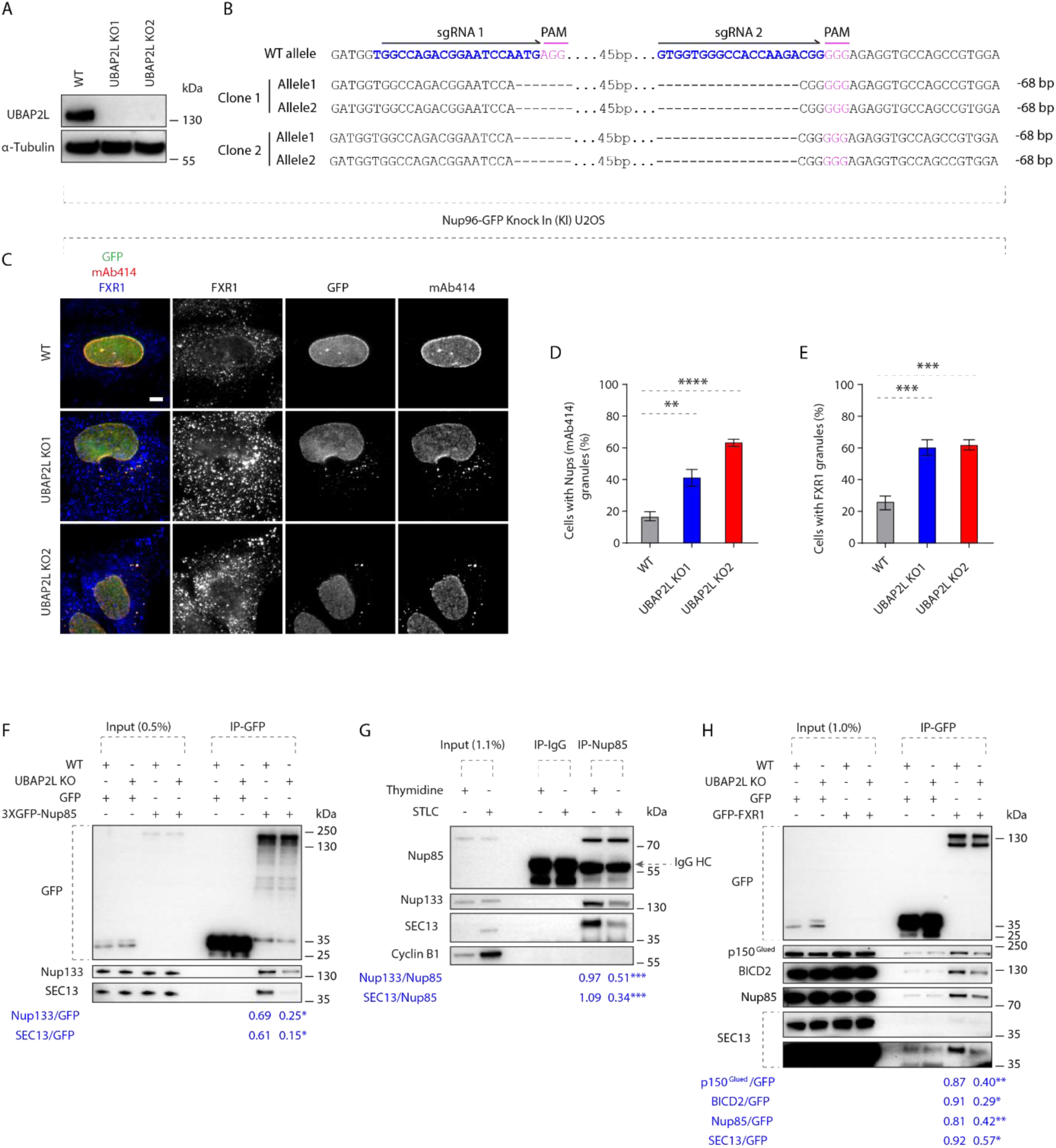
UBAP2L regulates the interaction between FXR1 and Y-complex Nups. (**A and B**) Validation of CRISPR/Cas9-mediated UBAP2L KO Nup96-GFP KI U2OS cell clones by Western blot (**A**) and Sanger sequencing (**B**). (**C to E**) Representative immunofluorescence images of the localization of Nups (GFP-Nup96 and mAb414) and FXR1 in WT and in two UBAP2L KO Nup96-GFP KI U2OS clonal cell lines in interphase cells synchronized by DTBR at 15h (**C**). Nuclei were stained with DAPI. The percentage of cells with cytoplasmic granules of Nups (mAb414) (**D**) and of FXR1 (**E**) shown in (**C**) were quantified. At least 200 cells per condition were analyzed (mean ± SD, **P < 0.01, ***P < 0.001, ****P < 0.0001, two-tailed t-test, *N* = 3). Scale bar, 5 μm. **(F)** Lysates of WT and UBAP2L KO Hela cells expressing GFP alone or 3XGFP-Nup85 for 27h and synchronized in G1/S phase by Thymidine 16h were immunoprecipitated using agarose GFP-Trap A beads (GFP-IP), analyzed by Western blot and signal intensities were quantified (shown a mean value, *P < 0.05; *N* = 3). **(G)** HeLa cells lysates of cells synchronized in interphase (Thymidine 16h) and of cells synchronized in mitosis (STLC 16h) were immunoprecipitated using Nup85 antibody or IgG, analyzed by Western blot and signal intensities were quantified (shown a mean value, ***P < 0.001; *N* = 3). **(H)** Lysates of interphase WT and UBAP2L KO HeLa cells expressing GFP alone or GFP-FXR1 for 27h were immunoprecipitated using agarose GFP-Trap A beads (GFP-IP), analyzed by Western blot and signal intensities were (shown a mean value, *P < 0.05, **P < 0.01; *N* = 3).

**Fig. S5.**
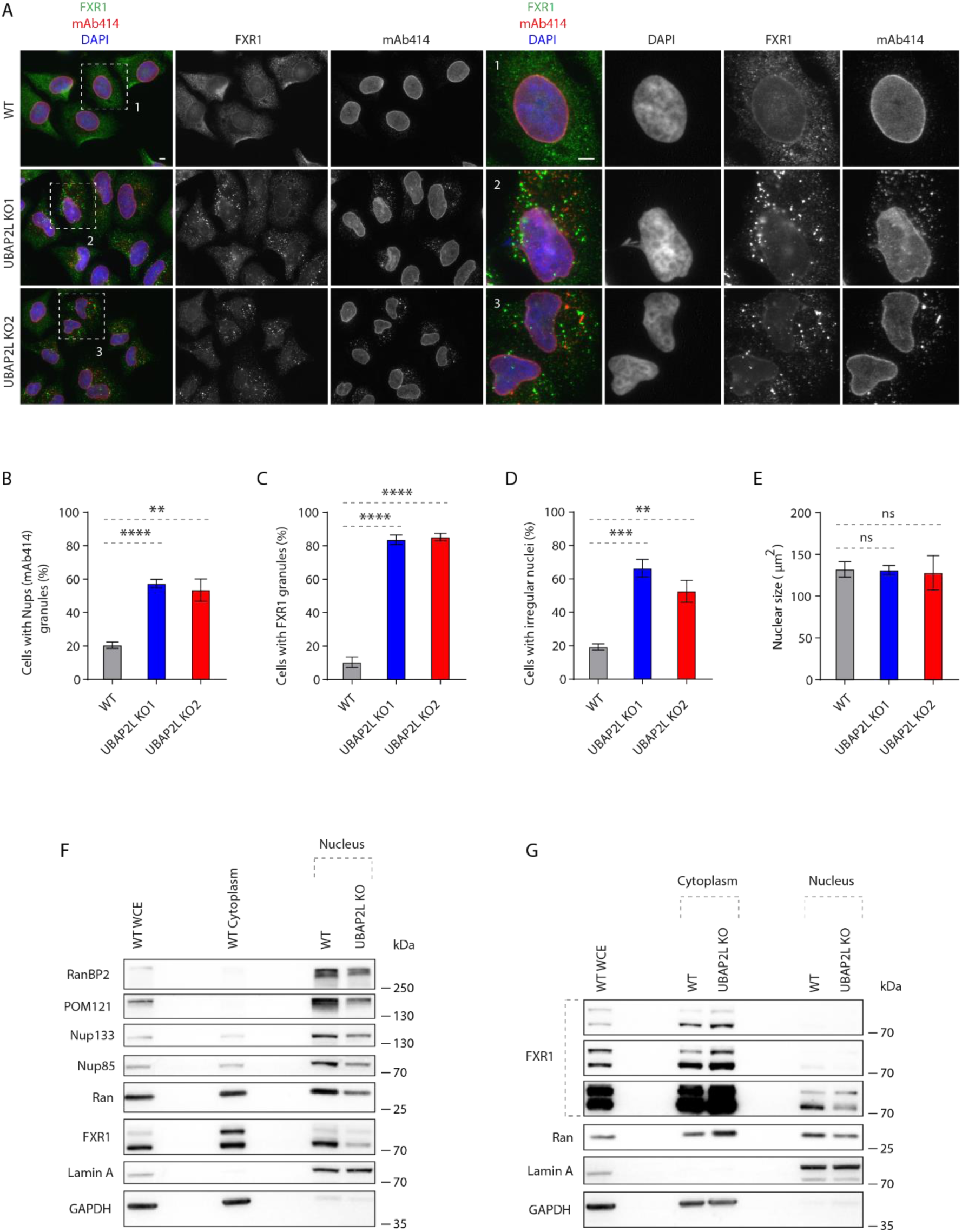
UBAP2L regulates localization of Nups and FXR1. (**A to E**) Representative immunofluorescence images depicting the nuclear shape and localization of Nups (mAb414) and FXR1 in WT and UBAP2L KO HeLa cells in interphase cells synchronized by DTBR at 12h (**A**). Nuclei were stained with DAPI. The percentage of cells with cytoplasmic granules of Nups (mAb414) (**B**) and of FXR1 (**C**) and irregular nuclei (**D**) and the nuclear size (**E**) shown in (**A**) were quantified. At least 250 cells per condition were analyzed (mean ± SD, ns, non-significant, **P < 0.01, ***P < 0.001, ****P < 0.0001, two-tailed t-test, *N* = 3). The magnified framed regions are shown in the corresponding numbered panels. Scale bars, 5 μm. The magnified framed regions are shown in the corresponding numbered panels. Scale bars, 5 μm. **(F)** The nuclear and cytoplasmic protein levels of Nups and NPC transport-associated factors in WT and UBAP2L KO HeLa cells synchronized as in (**A**) were analyzed by Western blot. WCE indicates whole cell extract. **(G)** The nuclear and cytoplasmic protein levels of Nups and NPC transport-associated factors in in asynchronously proliferating WT and UBAP2L KO HeLa cells were analyzed by Western blot. WCE indicates whole cell extract.

**Fig. S6.**
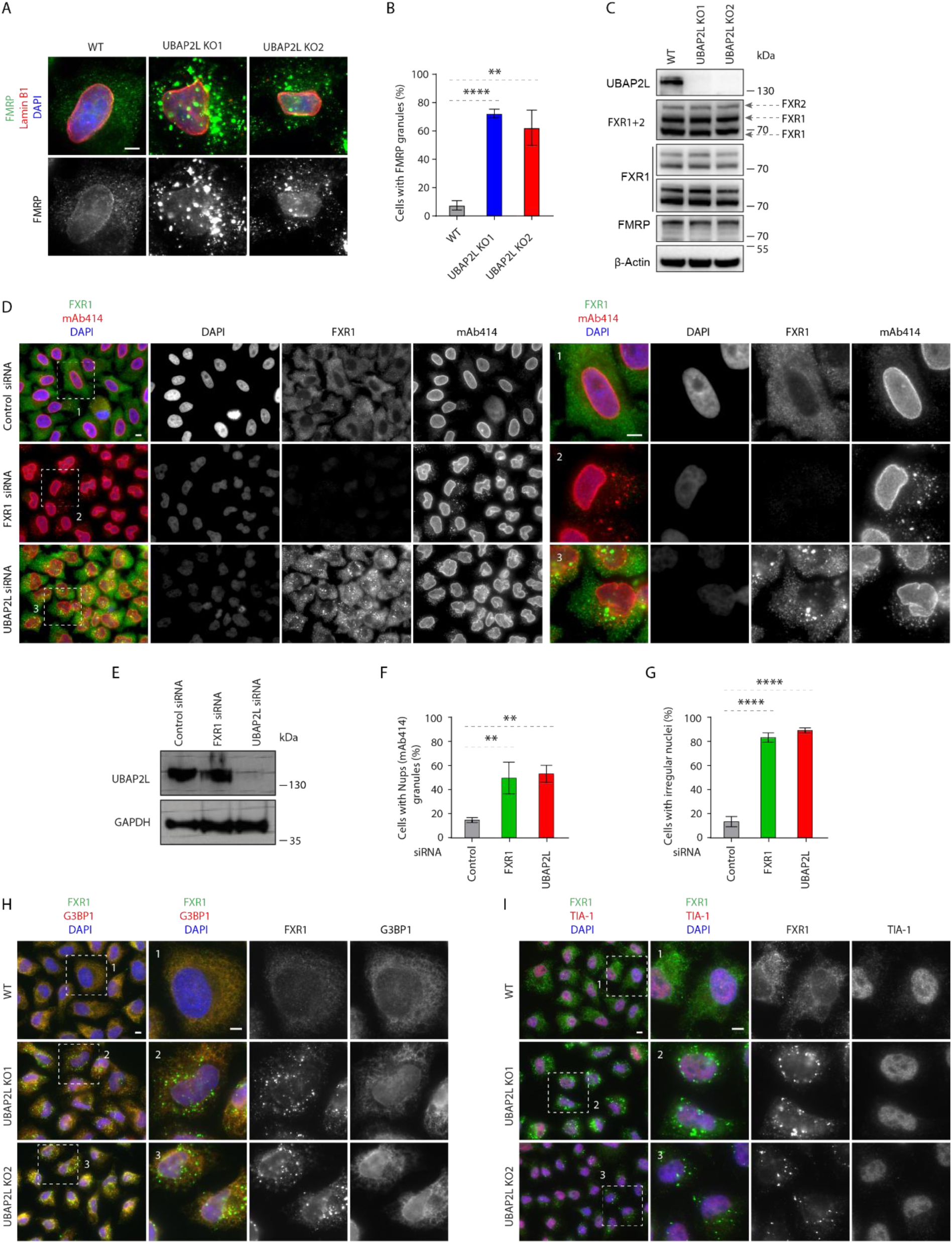
UBAP2L regulates FXRP proteins in the cytoplasm. (**A and B**) Representative immunofluorescence images depicting the localization of FMRP and Lamin B1 in WT and UBAP2L KO HeLa cells synchronized in interphase by DTBR at 12h (**A**). Nuclei were stained with DAPI. The percentage of cells with the cytoplasmic granules containing FMRP shown in (**A**) was quantified (**B**). At least 200 cells per condition were analyzed (mean ± SD, \*\**P* < 0.01, \*\*\*\**P* < 0.0001, two-tailed *t*-test, *N* = 3). Scale bar, 5 μm. (**C**) The protein levels of FXRP proteins in WT and UBAP2L KO HeLa cells synchronized in interphase by DTBR at 12h were analyzed by Western blot. (**D to G**) Representative immunofluorescence images depicting localization of FXR1 and Nups (mAb414) and the nuclear shape in the HeLa cells treated with indicated siRNAs and synchronized in interphase by DTBR at 12h (**D**). Nuclei were stained with DAPI. The magnified framed regions are shown in the corresponding numbered panels. UBAP2L protein levels in (**D**) were analyzed by Western blot (**E**). The percentage of cells with the cytoplasmic granules of Nups (mAb414) (**F**) and irregular nuclei (**G**) shown in (**D**) were quantified. At least 200 cells per condition were analyzed (mean ± SD, \*\**P* < 0.01, \*\*\*\**P* < 0.0001, two-tailed *t*-test, *N* = 3). Scale bars, 5 μm. (**H and I**) Representative immunofluorescence images of WT and UBAP2L KO HeLa cells synchronized in interphase by DTBR at 12h under non-stress conditions depicting localization of FXR1 (**H, I**), G3BP1 (**H**) and TIA-1 (**I**). Nuclei were stained with DAPI. Note that FXR1-containing granules present in non-stressed UBAP2L KO HeLa cells do not co-localize with stress granule (SG) components. The magnified framed regions are shown in the corresponding numbered panels. Scale bars, 5 μm.

**Fig. S7.**
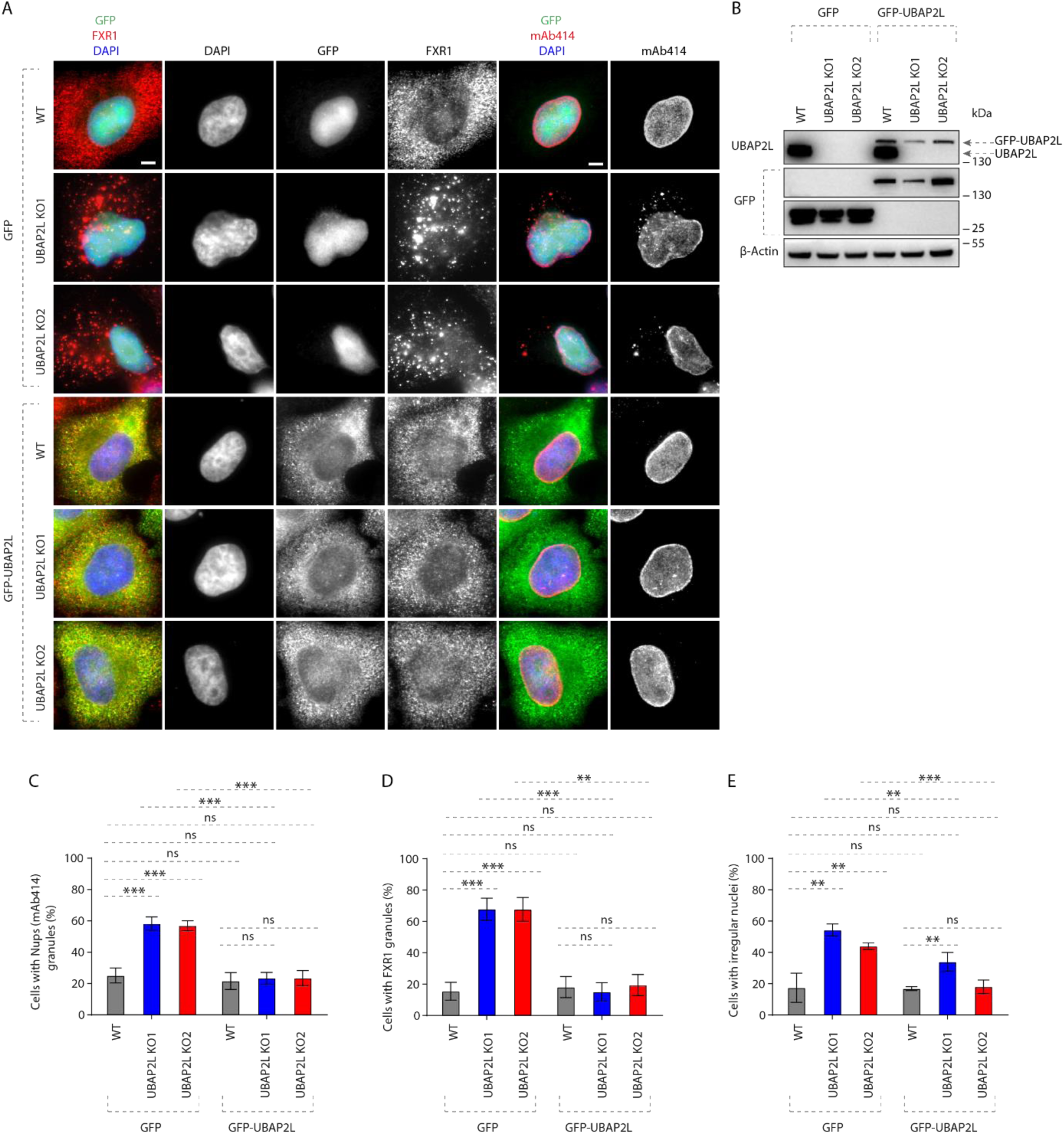
UBAP2L specifically regulates localization of Nups and FXR1 and nuclear shape. (**A and B**) Representative immunofluorescence images depicting the nuclear shape and localization of FXR1 and Nups (mAb414) in WT and UBAP2L KO HeLa cells expressing GFP alone or GFP-UBAP2L for 60h and synchronized in interphase by DTBR at 12h (**A**). Nuclei were stained with DAPI. Note that ectopic expression of GFP-UBAP2L but not GFP can rescue the nuclear and localization phenotypes in both UBAP2L KO HeLa cell lines. Scale bars, 5 μm. The protein levels of endogenous UBAP2L, GFP and GFP-UBAP2L of cells shown in (**A**) were analyzed by Western blot (**B**). (**C to E**) The percentage of cells with the cytoplasmic granules of Nups (mAb414) (**C**) and of FXR1 (**D**) and irregular nuclei (**E**) shown in (**A**) were quantified. At least 200 cells per condition were analyzed (mean ± SD, ns: not significant, **P < 0.01, ***P < 0.001, two-tailed *t*-test, *N* = 3).

**Fig. S8.**
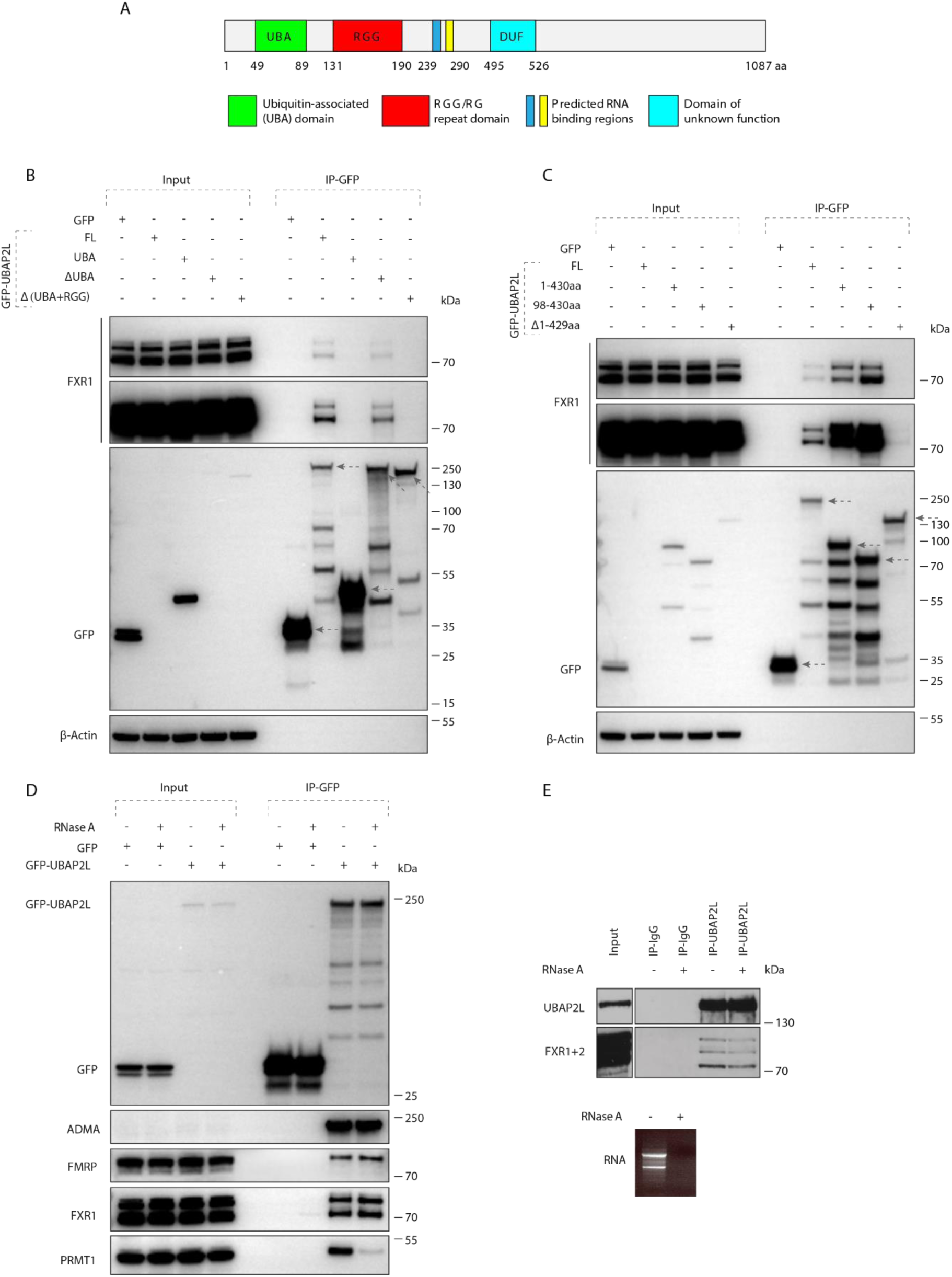
Arginines within the RGG domain of UBAP2L are required for the interaction with FXR1. **(A)** Domain organization of UBAP2L depicting UBA domain, RGG/RG repeat domain, two predicted RNA binding regions and the domain of unknown function (DUF). (**B and C**) Lysates of HeLa cells expressing GFP alone or GFP-UBAP2L-dervied constructs (full length FL, UBA, ΔUBA or Δ(UBA+RGG) fragments) for 27h were immunoprecipitated using agarose GFP-Trap A beads (GFP-IP) and analyzed by Western blot (**B**). Lysates of HeLa cells expressing GFP alone or several GFP-UBAP2L-dervied constructs (FL, 1-430 aa, 98-430 aa or Δ1-429 aa fragments) for 27h were immunoprecipitated using agarose GFP-Trap A beads (GFP-IP) and analyzed by Western blot (**C**). The arrows indicate the bands corresponding to the expressed GFP proteins while the remaining bands are non-specific. (**D and E**) Interphase HeLa cells expressing GFP alone or GFP-UBAP2L for 27h and cell lysates were treated with RNase A, immunoprecipitated using agarose GFP-Trap A beads (GFP-IP) and analyzed by Western blot. Note that RNase treatment can abolish interaction with PRMT1 but not with FXRPs (**D**). Immunoprecipitations from cell lysates of HeLa cells treated with RNase A using UBAP2L antibody or IgG were analyzed by Western blot. Efficiency of the RNase treatment was confirmed by imaging of mRNAs by agarose gel electrophoresis and ethidium bromide staining (**E**).

**Fig. S9.**
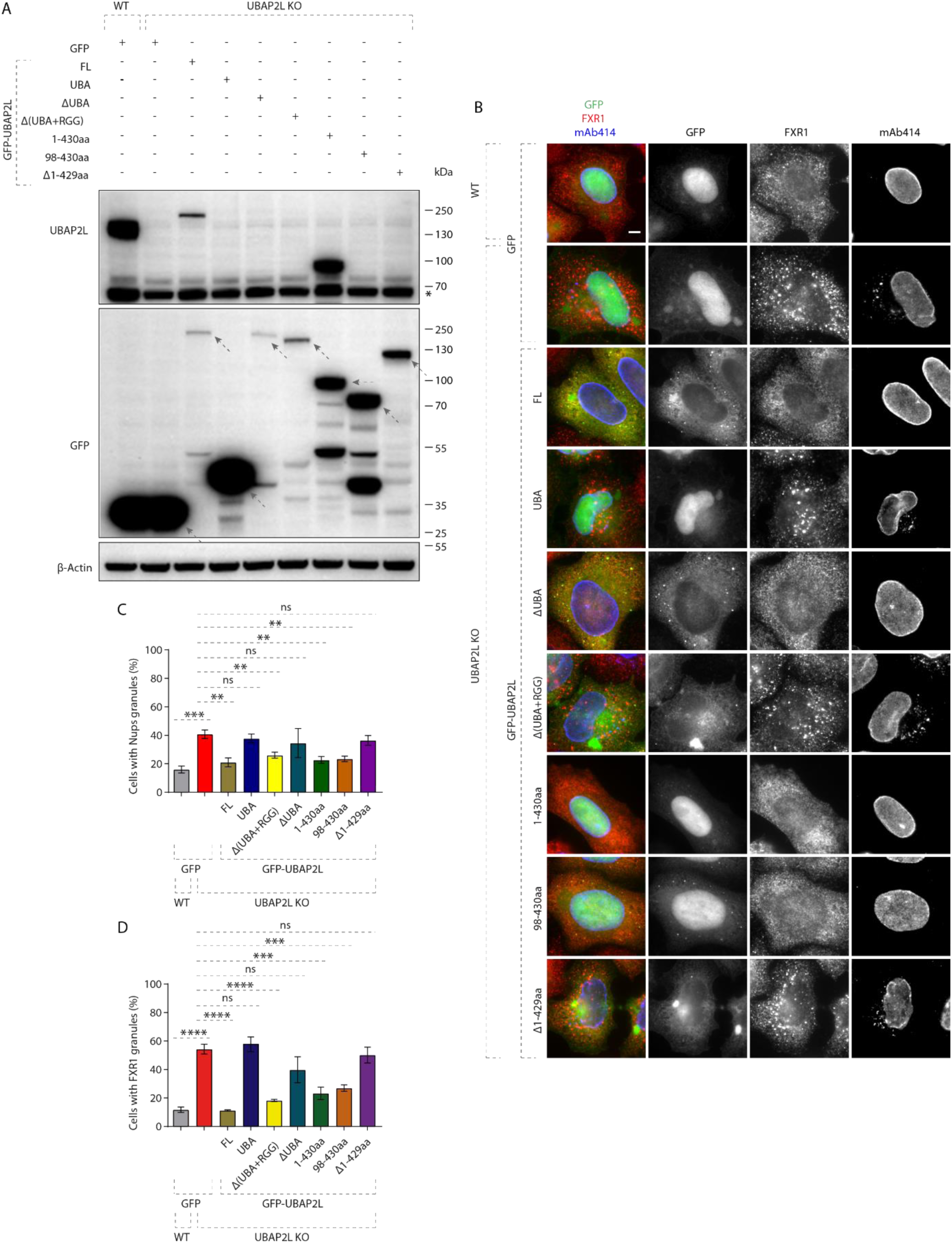
98-430 aa fragment of UBAP2L protein is required for the the function of UBAP2L on Nups and FXR1. **(A)** The protein levels of endogenous UBAP2L, GFP and GFP-UBAP2L-derived versions (FL, UBA, ΔUBA, Δ(UBA+RGG), 1-430 aa, 98-430 aa or Δ1-429 aa) of cells shown in (B) were analyzed by Western blot. The arrows indicate the bands corresponding to the expressed GFP proteins while the remaining faster migrating bands are either non-specific or degradation products. (**B to D**) Representative immunofluorescence images depicting localization of FXR1 and Nups (mAb414) in WT and UBAP2L KO HeLa cells expressing GFP alone or GFP-UBAP2L-derived fragments (FL, UBA, ΔUBA, Δ(UBA+RGG), 1-430 aa, 98-430 aa or Δ1-429 aa) for 60h and synchronized in interphase by DTBR at 12h (**B**). Note that the UBAP2L 98-430 aa protein fragment containing the RGG domain is required for the function of UBAP2L on Nups. The percentage of cells with the cytoplasmic granules of Nups (mAb414) (**C**) and of FXR1 (**D**) shown in (**B**) were quantified. At least 200 cells per condition were analyzed (mean ± SD, ns: not significant, **P < 0.01, ***P < 0.001, ****P < 0.0001, two-tailed *t*-test, *N* = 3). Scale bar, 5 μm.

**Fig. S10.**
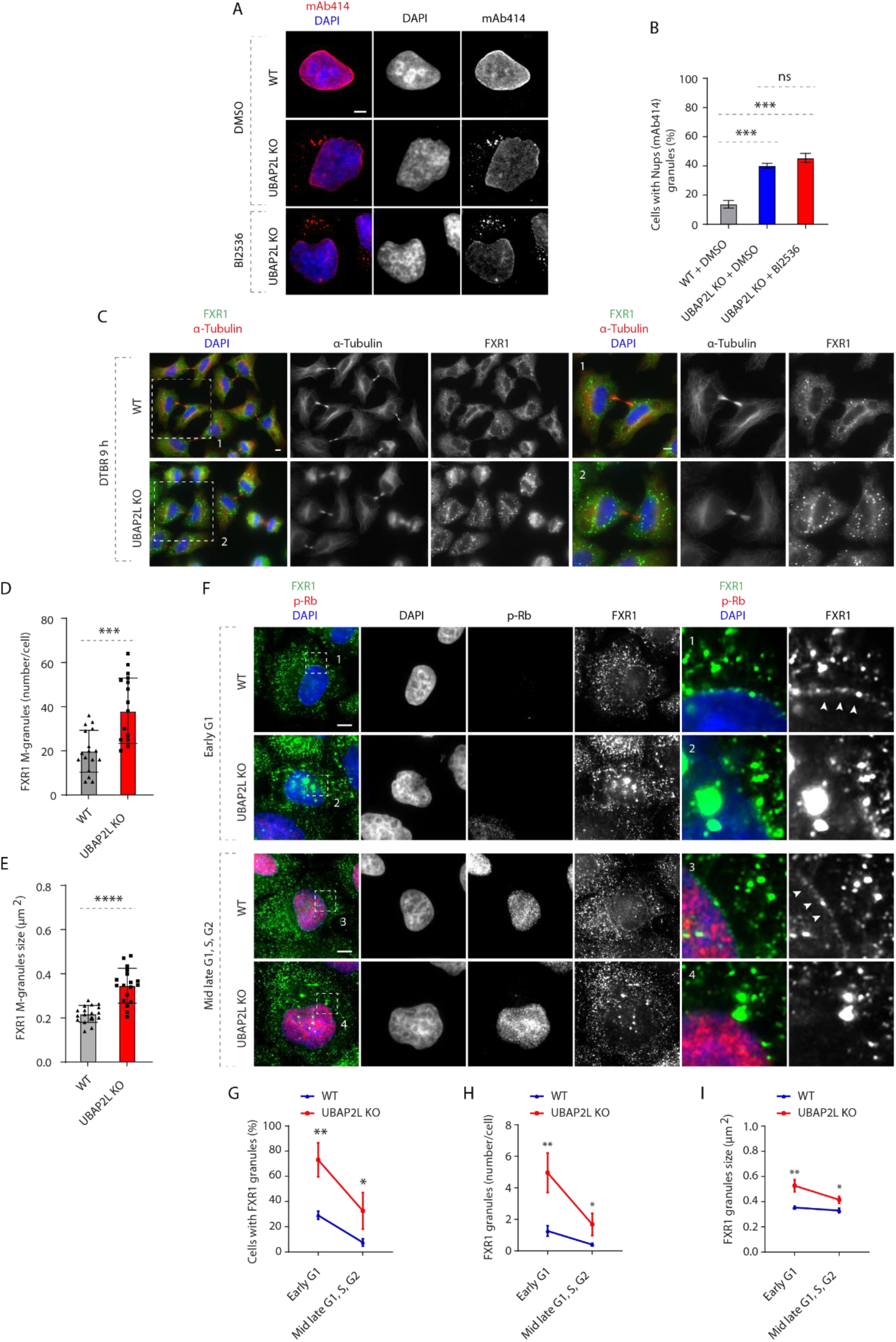
UBAP2L drives localization of FXR1 to the NE during early G1. (**A and B**) Representative immunofluorescence images depicting the localization of Nups (mAb414) in WT and UBAP2L KO HeLa cells synchronized in interphase by double thymidine block and release (DTBR) at 12h (**A**). PLK1 inhibitor BI 2536 (or solvent control) was used at a concentration of 100 nM for 45 min prior to sample collection. Nuclei were stained with DAPI. The percentage of cells with the cytoplasmic granules containing Nups (mAb414) shown in (**A**) was quantified (**B**). At least 150 cells per condition were analyzed (mean ± SD, ns: not significant, ***P < 0.001, two-tailed t-test, *N* = 3). Scale bar, 5 μm. (**C to E**) Representative immunofluorescence images depicting the localization of FXR1 in WT and UBAP2L KO HeLa cells synchronized by DTBR 9h in late telophase (**C**). Nuclei were stained with DAPI. The magnified framed regions are shown in the corresponding numbered panels. Scale bars, 5 μm. The number of FXR1 granule per cell (number/cell) (**D**) and the size of FXR1 granules (granule ≥ 0.105 µm^2^) (**E**) shown in (**C**) were quantified. 17 WT and 18 UBAP2L KO HeLa cells were counted, respectively. (**F to I**) Representative immunofluorescence images depicting the localization of FXR1 in different cell cycle stages in asynchronously proliferating WT and UBAP2L KO HeLa cells (**F**). p-Rb was used to distinguish between early G1 (p-Rb-negative cells) and mid-late G1, S and G2 (p-Rb-positive cells) stages. Nuclei were stained with DAPI. The arrowheads indicate the nuclear envelope (NE) localization of endogenous FXR1. Scale bars, 5 μm. The percentage of cells with the cytoplasmic FXR1 granules (**G**), the number of FXR1 granule per cell (number/cell) (**H**) and the size of FXR1 granules (granule ≥ 0.2109 µm^2^) (**I**) shown in (**F**) were quantified. At least 200 cells per condition were analyzed (mean ± SD, ns: not significant, *P < 0.05, **P < 0.01, two-tailed t-test, *N* = 3).

**Fig. S11.**
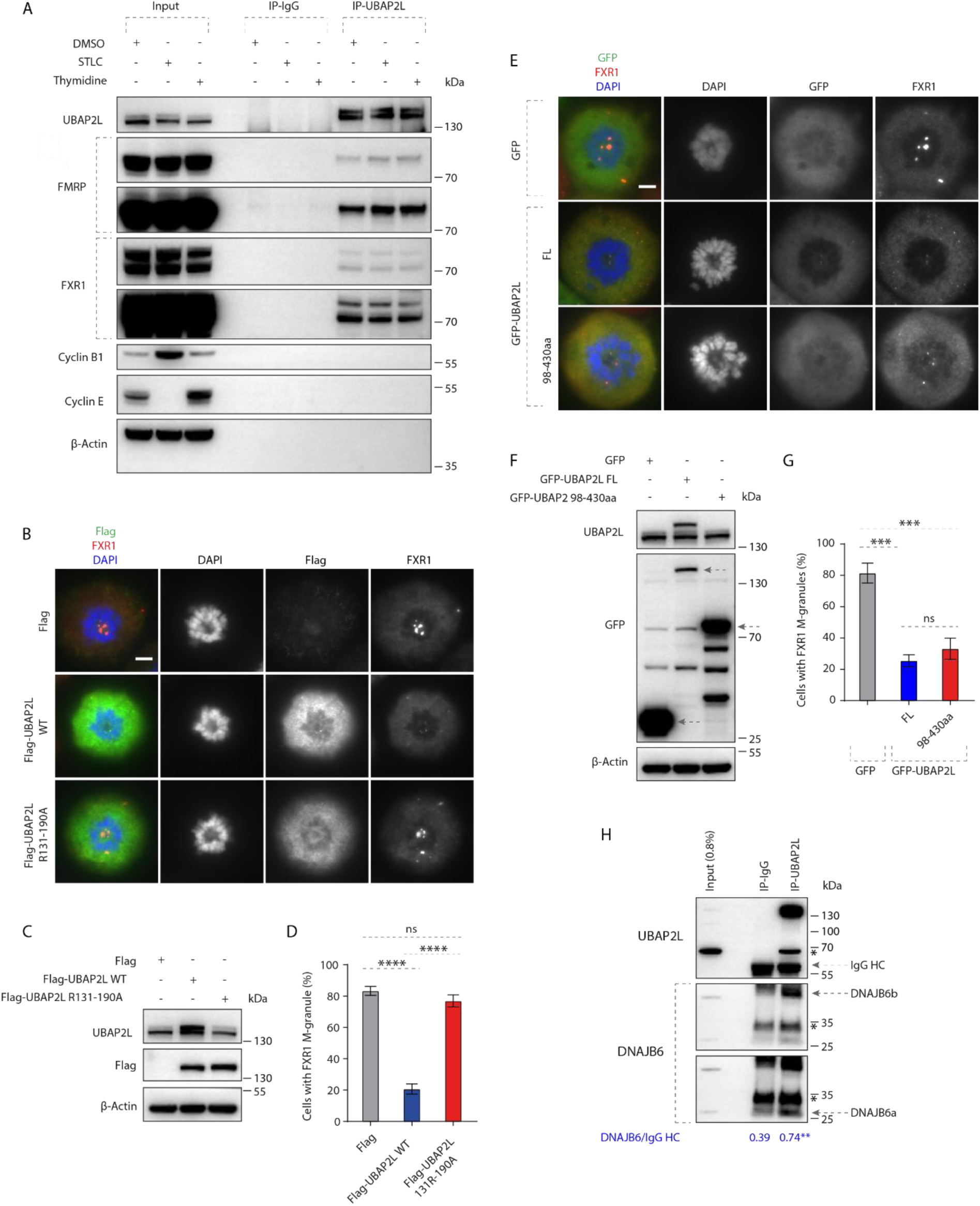
UBAP2L can dissolve FXR1-containing mitotic foci. **(A)** Immunoprecipitations from HeLa cells lysates of asynchronously proliferating cells (DMSO 16h), cells synchronized in mitosis (STLC 16h) or in interphase (thymidine 16h) using UBAP2L antibody or IgG were analyzed by Western blot. (**B to D**) HeLa cells expressing Flag, Flag-UBAP2L WT or Flag-UBAP2L R131-190A for 27h were synchronized in prometaphase using STCL for 16h and representative immunofluorescence images depicting localization of FXR1 are shown in (**B**). Chromosomes were stained with DAPI. The protein levels of Flag-UBAP2L and endogenous UBAP2L in (**B**) were analyzed by Western blot (**C**). The percentage of cells with FXR1-granules shown in (**B**) were quantified (**D**). At least 200 cells per condition were analyzed (mean ± SD, ns: not significant, ****P < 0.0001, two-tailed *t*-test, *N* = 3). Scale bar, 5 μm. (**E to G**) Representative immunofluorescence images depicting the localization of FXR1 in HeLa cells expressing GFP, GFP-UBAP2L FL or GFP-UBAP2L 98-430aa for 27h synchronized in prometaphase using STCL for 16h (**E**). Chromosomes were stained with DAPI. The protein levels of GFP-UBAP2L and endogenous UBAP2L in (**E**) were analyzed by Western blot (**F**). The percentage of cells with FXR1-granules shown in (**E**) was quantified (**G**). At least 200 cells per condition were analyzed (mean ± SD, ns: not significant, ***P < 0.001, two-tailed *t*-test, *N* = 3). Scale bar, 5 μm. (**H**) HeLa cells lysates were immunoprecipitated from using UBAP2L antibody or IgG, analyzed by Western blot and signal intensities were quantified (shown a mean value, **P < 0.01; *N* = 3). The arrows indicate the bands corresponding to the IgG heavy chain (HC) and to DNAJB6a and b, respectively . * Indicates non-specific bands.

**Fig. S12.**
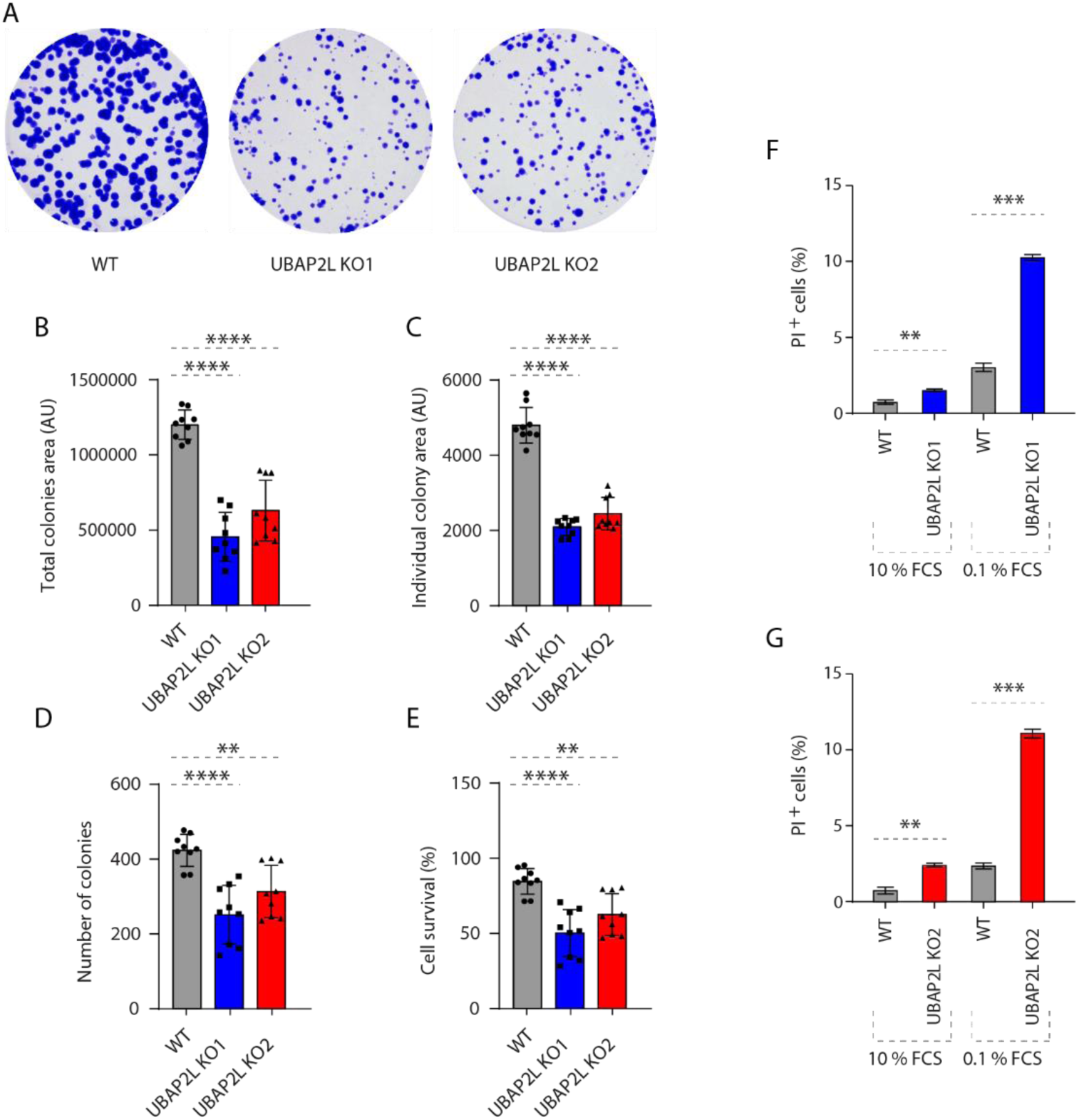
UBAP2L regulates long-term proliferation capacity of HeLa cells and ensures survival of HeLa cells upon nutrient stress. (**A to E**) Representative images of colony formation assays of WT and UBAP2L KO HeLa cells maintained in culture for 7 days (**A**). Total colony area (**B**), individual colony area (**C**), average number of colonies (**D**) and cell survival (**E**) of cells shown in (**A**) were quantified using the Fiji software (mean ± SD, *P < 0.05, **P < 0.01, ***P < 0.001, ****P < 0.0001; two-tailed t-test, *N* = 3). (**F and G**) The percentage of propidium Iodide (PI)-positive cells in WT and UBAP2L KO HeLa cells cultured in the indicated concentrations of serum for 72h were quantified by fluorescence activated cell sorting (FACS) (mean ± SD, **P < 0.01, ***P < 0.001, two-tailed t-test, *N* = 3).

### Supplementary tables

**Table S1.**
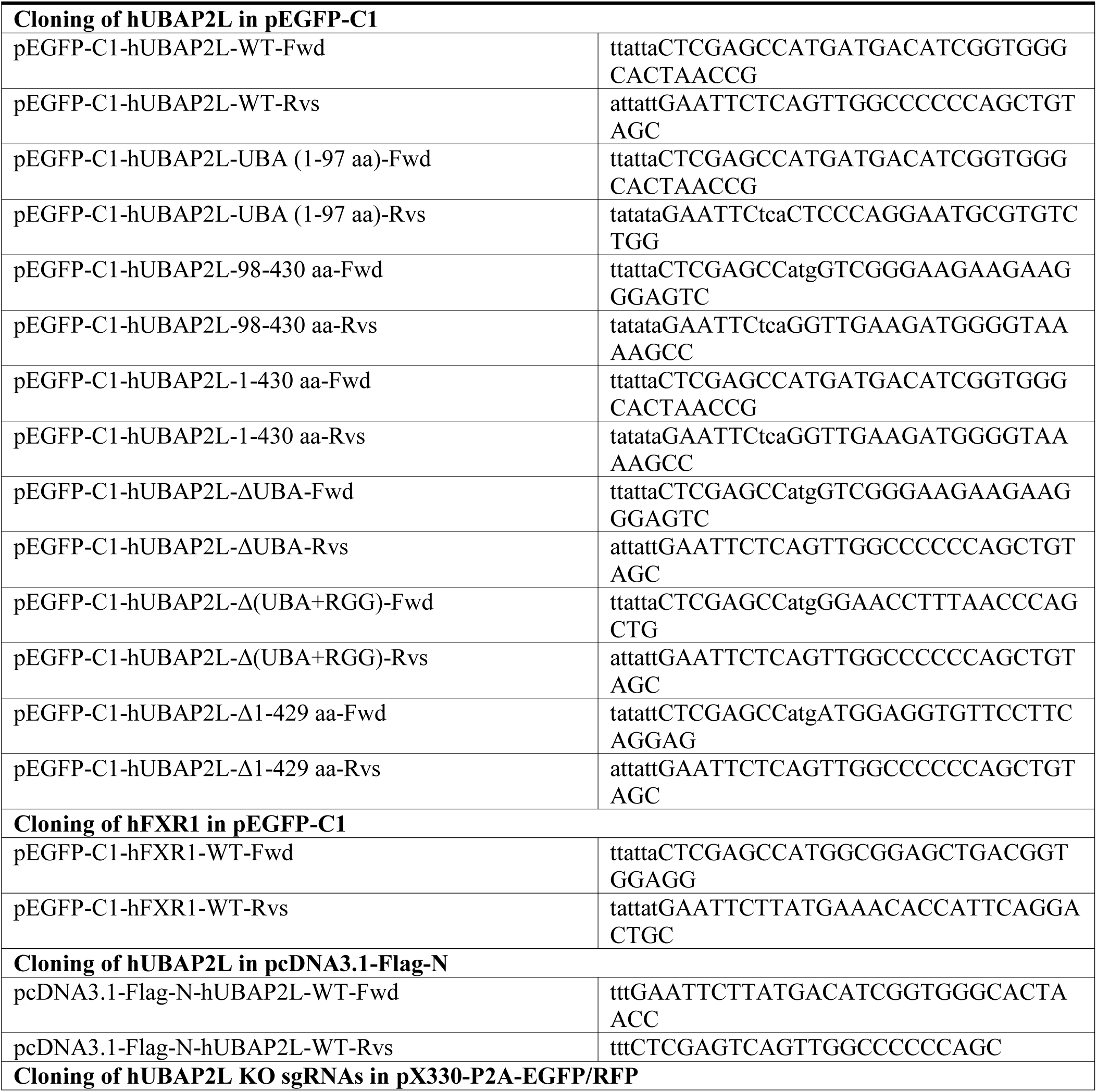
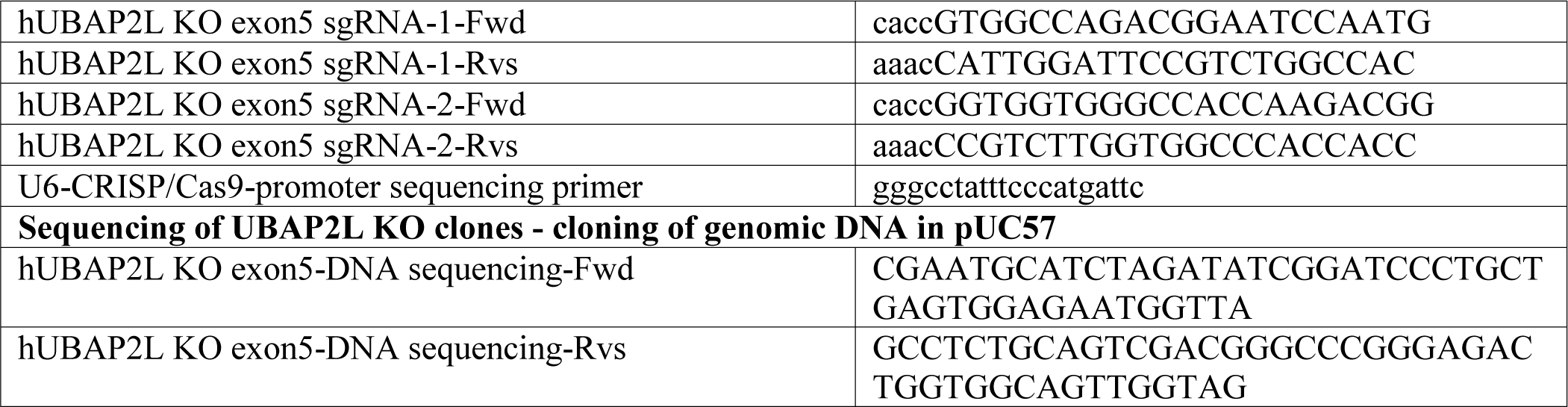
Cloning primers. describes the cloning primers used in the study

**Table S2.** Reagents and resources. describes other reagents and resources including bacterial stains, cell lines, chemicals, cDNAs and software used in the study

| REAGENT or RESOURCE | SOURCE | IDENTIFIER |
| --- | --- | --- |
| **Bacterial strains** |  |  |
| DH5alpha Competent *E. coli* | NEW ENGLAND BioLabs | Cat# C2987I |
| **Chemicals and Peptides** |  |  |
| Thymidine | Sigma-Aldrich | Cat# T1895-5G |
| Nocodazole | Sigma-Aldrich | Cat# M-1404 |
| Monastrol | Sigma-Aldrich | Cat# M8515 |
| 4',6-Diamidino-2-phenylindole dihydrochloride (DAPI) | Sigma-Aldrich | Cat# D8417 |
| MG132 | Tocris Bioscience | Cat# 1748 |
| STLC (S-Trityl-L-cysteine) | Enzo Life Sciences | Cat# ALX-105-011-M500 |
| MOWIOL 4-88 Reagent | Millipore | Cat# **475904-M** |
| jetPEI®-DNA transfection reagent | Polyplus transfection | Cat# 101-01N |
| SiR-DNA | Spirochrom | Cat# SC007 |
| Lipofectamine™ 2000 Transfection Reagent | Invitrogen | Cat# 11668019 |
| Lipofectamine™ RNAiMAX Transfection Reagent | Invitrogen | Cat# 13778150 |
| X-tremeGENE™ 9 DNA Transfection Reagent | Roche | Cat# 6365787001 |
| Dexamethasone | Sigma-Aldrich | Cat# D8833 |
| T4 DNA Ligase | New England Biolabs | Cat# M0202T |
| Exonuclease III | Takara | Cat# 2170B |
| Cycloheximide | Sigma-Aldrich | Cat# C4859 |
| Glucose oxidase | Sigma-Aldrich | Cat# G2133 |
| Cyclooctatetraene | Sigma-Aldrich | Cat# 138924 |
| Catalase | Sigma-Aldrich | Cat# C1345 |
| Leptomycin B | Abcam | Cat# ab120501 |
| Lovastatin | Sigma-Aldrich | Cat# 75330-75-5 |
| Psoralidin | Sigma-Aldrich | Cat# 18642-23-4 |
| Benzonase® Nuclease | Millipore | Cat# 70746 |
| RNAse A | Thermo Fisher Scientific | Cat# EN0531 |
| Propidium iodide (PI) | Sigma-Aldrich | Cat# P4170 |
| Phalloidin 488 | Thermo Fisher Scientific | Cat# A12379 |
| **Cell Lines** |  |  |
| Human: HeLa (Kyoto) | ATCC | Cat# CCL-2 |
| Human: HeLa UBAP2L KO | This study | N/A |
| Human: U2OS bone osteosarcoma | ATCC | Cat# HTB-96 |
| Human: Nup96-GFP KI U2OS | Arnaud Poterszman (IGBMC) | N/A |
| Human: Nup96-GFP KI U2OS UBAP2L KO | This study | N/A |
| **Oligonucleotides** |  |  |
| siRNA: Non-targeting siGENOME | Dharmacon | Cat# D-001210-02-05 |
| siRNA: FXR1 individual | Dharmacon | Cat# J-012011-06-0005 |
| siRNA: UBAP2L individual | Dharmacon | Cat# J-021220-09-0002 |
| Primers used for Cloning and Sequencing are described in Table S1 | This study | N/A |
| **Recombinant DNA** |  |  |
| pcDNA3.1-Flag-N | This study | N/A |
| pcDNA3.1-Flag-N-UBAP2L WT | This study | N/A |
| pcDNA3.1-Flag-UBAP2L R131-190A | (Huang *et al*, 2020) | N/A |
| pEGFP-C1 | Clontech | Cat# 6084-1 |
| pEGFP-C1-UBAP2L WT | This study | N/A |
| pEGFP-C1-UBAP2L UBA | This study | N/A |
| pEGFP-C1-UBAP2L ΔUBA | This study | N/A |
| pEGFP-C1-UBAP2L 98-430aa | This study | N/A |
| pEGFP-C1-UBAP2L 1-430aa | This study | N/A |
| pEGFP-C1-UBAP2L Δ1-429aa | This study | N/A |
| pEGFP-C1-UBAP2L Δ(ΔUBA+RGG) | This study | N/A |
| pEGFP-C1-FXR1 WT | This study | N/A |
| pEGFP-C1-Nup85 WT | Valérie Doye (Institut Jacques Monod, Paris) | N/A |
| pXRGG-GFP | (Hamada *et al*, 2011; Love *et al*, 1998) | N/A |
| pUC57 | Thermo | Cat# SD0171 |
| pX330-P2A-EGFP | (Zhang *et al*, 2017) | N/A |
| pX330-P2A-RFP | (Zhang *et al*, 2017) | N/A |
| **Software and Algorithms** |  |  |
| CRISPR/Cas9 Guide RNA Design | Benchling | https://www.benchling.com/ |
| Fiji Image Analysis | ImageJ | https://imagej.net/Fiji |
| Prism | GraphPad | N/A |
| Illustrator | Adobe | N/A |
| MATLAB | Mathworks | N/A |

